# Selector: A General Python Library for Diverse Subset Selection

**DOI:** 10.1101/2025.11.21.689756

**Authors:** Fanwang Meng, Marco Martínez González, Valerii Chuiko, Alireza Tehrani, Abdul Rahman Al Nabulsi, Abigail Broscius, Hasan Khaleel, Kenneth López-Pérez, Ramón Alain Miranda-Quintana, Paul W. Ayers, Farnaz Heidar-Zadeh

## Abstract

`Selector` is a free, open-source Python library for selecting diverse subsets from any dataset, making it a versatile tool across a wide range of application domains. `Selector` implements different subset sampling algorithms based on sample distance, similarity, and spatial partitioning, along with metrics to quantify subset diversity. It is flexible and integrates seamlessly with popular Python libraries like `Scikit-Learn`, demonstrating the interoperability of the implemented algorithms with data analysis workflows. `Selector` is an operating-system agnostic, accessible, and easily extensible package designed with modern software development practices, including version control, unit testing, and continuous integration. Interactive quick-start notebooks, which are also web-accessible, provide user-friendly tutorials for all skill levels, showcasing applications in computational chemistry, drug discovery, and chemical library design. Additionally, a web interface has been developed that allows users to easily upload datasets, configure sampling settings, and run subset selection algorithms, with no programming required. This paper serves as the official release note for the `Selector` package, offering a technical overview of its features, use cases, and development practices that ensure its quality and maintainability.

## 1 Introduction

Sampling is a widely used approach for selecting representative subsets from larger datasets, particularly when collecting or analyzing every instance is impractical or resource-intensive. Poor sampling is known to introduce bias, compromise the validity of results, and lead to misleading conclusions. To mitigate this, selecting a diverse subset (commonly known as the diversity selection problem) is critical for data-driven modeling across various fields, including ecology^1,2^, geography^3^, computational chemistry^4–6^, computational biology^7^, drug discovery^8–10^, experimental design^11^, and machine learning (ML)^12,13^. For example, this is routinely used in high-throughput screening to identify di-versity voids and enrich chemical space.^14^

Identifying chemically diverse compounds, particularly those at the extremes of chemical space, is central to navigation tools like ChemGPS and ChemMaps.^15–17^ Diversity-based selection also mitigates biases such as class imbalance in ML training sets,^18–20^ facilitating the discovery of drug-like compounds and functional materials.^21^ Incorporating both structural and property data during sampling can reveal activity cliffs and ridges,^22,23^, which improves the performance and generalizability of ML models.^24^ Diverse sampling is also crucial for modeling the potential energy surface and reaction dynamics. Compared to enhanced sampling techniques such as weighted ensemble and replica exchange molecular dynamics (MD),^25^ diversity-based selection captures rare conformations more efficiently. This, in turn, accelerates simulation convergence^26^ and enables a more comprehensive exploration of the free-energy landscape.

Diverse exploration is also central to recent advances in inverse molecular design.^21,27,28^ Regardless of whether real-valued embeddings^29^ or string-based representations^30^ are used, the ability to navigate chemical space with minimal data remains a central challenge. This has driven the adoption of diversity-driven chemical library design^31,32^ and *de novo* molecular generation^33,34^. Diversity methods help uncover alchemical paths, indicating how structural mutations lead to unexplored chemical space.^35^ These methods have also improved sampling efficiency in reinforcement learning, demonstrating that agents rewarded for exploring diverse compounds are more successful at finding high-performance molecules.^36^

Despite the significance and broad applicability of sampling, there is currently no free, open-source, and user-friendly library dedicated to diversity-based subset selection algorithms and diversity measures. Many existing tools are limited by one or a few hardcoded algorithms and provide little to no support for diversity-based sampling or customizable similarity and distance measures. As a result, they often lack the flexibility and extensibility required for broader applications. For example, `DISSIM`^37^, a Fortran 77 implementation of the MaxMin algorithm from 1998^38^, was later reimplemented as the Windows-based `DivCalc`^39^ in 2002. The open-source `caret` package^40^ implements the MaxMin algorithm in R, while `DiversityNet` offers generative models for molecular diversity but lacks subset selection functionality. `PKOM`^41^ implements clustering algorithms for subset selection with no support for diversity-based sampling, on the other hand, `SubMo-GNN`^42^ uses graph neural networks to select diverse molecules based on measures like log-determinant functions and the Wasserstein distance to uniform distribution (WDUD). Similarly, `astartes`^43^ supports sphere exclusion^44^, Kennard-Stone sampling^45^, and OptiSim^46^ algorithms for ML dataset splits. More recently, tools such as `chemfp`^47^ and RDKit’s `SimDivFilters.rdSimDivPickers` module (as of version 2025.03.3)^48^ include methods like sphere exclusion^44^, MaxMin^38^, hierarchical clustering^49^, and Leader algorithm; however, they do not offer formal diversity metrics or an accessible framework for implementing other sampling algorithms. In contrast, `EvolMol`^50^ incorporated Shannon entropy to assess the diversity of *de novo* generated molecules, but does not include sampling algorithms.

In addition, these tools rarely use modern software development workflows such as unit testing, detailed documentation, continuous integration, and deployment (CI/CD), which limits their long-term usability and maintainability. This highlights the need for a broadly applicable, flexible, and user-friendly library that provides robust implementations of subset selection algorithms and diversity measures. `Selector` bridges this gap by providing a free, open-source, and cross-platform Python package that follows best practices in sustainable scientific software development. It also stands out as the only library offering a comprehensive and growing suite of diversity sampling algorithms and diversity metrics. `Selector` is designed with an intuitive and general-purpose Application Programming Interface (API) that supports a broad range of scientific domains. Here, we present an overview of `Selector`’s design principles, scope, and supported methods, and showcase its applications across various domains.

## 2 About Selector

`Selector` is a free, open-source library hosted on GitHub, distributed under the GNU General Public License, and maintained as part of the QC-Devs International Software Consortium which supports a broad range of computational packages for quantum chemistry^51–61^ and data science^62,63^. Following best practices in scientific software development^64^, we provide comprehensive documentation that includes algorithms and examples to support ease of use and extensibility. `Selector` is built on standard scientific Python libraries such as `NumPy`^65^, and maintains high code quality through a comprehensive suite of quality-assurance tools, including `black`, `pylint`, `autopep8`, `pycodestyle`, `bandit`, and `flake8`. Unit testing is handled with `pytest`^66^ and coverage is tracked via `codecov.io`, ensuring the robustness of the code during updates, releases, and installation. The package currently maintains 95% test coverage.

Continuous integration via GitHub Actions automates the execution of quality-assurance checks for every code change, making it a central part of our code review workflow. Continuous delivery enables us to build, test, and distribute `Selector` via the `pip` package manager. This lets users install the library and its dependencies with a single command. To access the latest features before an official release, users can install `Selector` directly from the development branch of the source code. To avoid outdated instructions, we do not include static installation steps in this paper and instead refer users to the `Selector` website for the latest installation procedures, documentation, and feature updates. We also offer interactive Jupyter notebook tutorials, which can be accessed via the web, to demonstrate the functionality of the package. In addition, the `Selector` web interface allows one to use the package directly through their browser, without writing code.

### 2.1 Architecture and Scope

Figure 1 sketches two key modules of this library. The `methods` module implements customizable algorithms for diverse selection based on distance, similarity, or partitioning of the feature space. It supports stratified subset selection by allowing users to provide sample labels, in which case, the algorithm automatically selects samples from each cluster. The `measures` module contains functions to quantify the diversity of subsets, calculate similarity, and convert similarity measures to distances.

**Figure 1.**
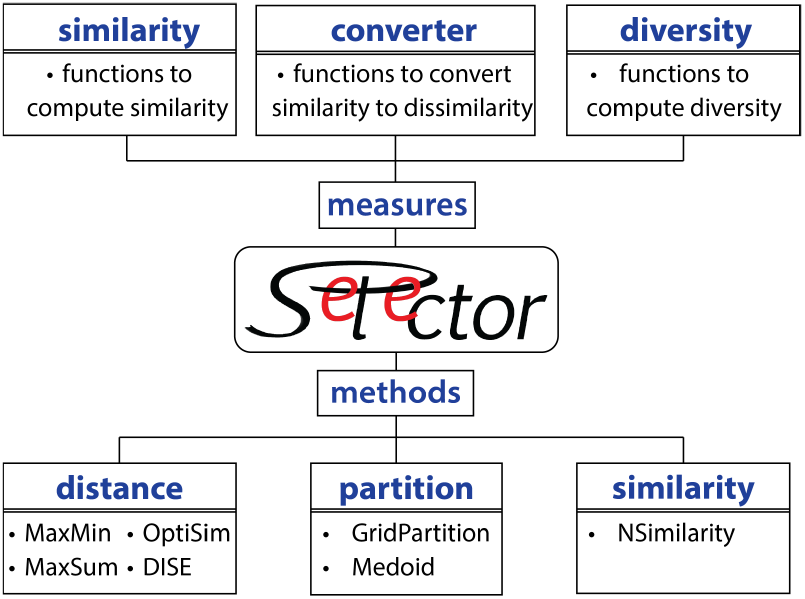
Schematic representation of key modules of the `Selector` package.

An often overlooked yet critical aspect of diverse subset selection is the choice of feature representation and pairwise distance function. Since these choices are highly context-dependent and vary with the specific problem, their selection and optimization are beyond the scope of this package. In this regard, `Selector` is designed to be agnostic to feature types and supports any user-defined distance metric. This demonstrates the interoperability of the subset selection algorithms implemented.

### 2.2 Subset Selection Methods

Here, we present an overview of the implemented algorithms for diverse subset selection, along with an analysis of their computational costs and suitability for different use cases. Figure SI 1 compares the supported methods in two dimensions.

#### 2.2.1 Distance-Based Selection Algorithms

Distance-based selection algorithms aim to maximize dissimilarity between samples by considering only pairwise distances. They are applicable in both high- and low-dimensional settings, but they are limited in addressing tasks like identifying diversity voids.^14^

The MaxMin^38^ and MaxSum^67^ both (1) start by selecting the initial point as the mediod, or a user-specified data point(s), (2) compute distance or dissimilarity of the available samples to the previously selected one(s) given a user-defined distance function, (3) select the next sample following a specific selection rule, and (4) iterate until the required number of samples is selected. In each iteration, the MaxMin method maximizes the minimum distance to the selected samples, whereas the MaxSum method maximizes the sum of distances to the selected samples. As both algorithms require the full pairwise distance matrix, their computational cost is 𝒪(*n*^2^*s*)^68^, where *s* is the total number of data points and *d* is the number of dimensions or features representing each data.

The OptiSim^46^ and Directed Sphere Exclusion (DISE)^69^ algorithms exclude all neighboring samples within a predetermined radius *r* (referred to as the exclusion sphere) around the selected sample and optimize the radius *r* to achieve a subset size that closely matches the desired target size, *n*, specified by the user. To select samples, OptiSim (1) starts with a reference sample and excludes all neighboring samples within radius *r*, and (2) iteratively chooses a random subsample of size *k*, and selects the sample with the greatest minimum Minkowski *p*-norm distance to the previously selected samples. On the other hand, DISE (1) sorts all samples in ascending order based on their Minkowski *p*-norm distance from a reference sample and excludes all samples within the radius *r* of the reference sample, (2) iterating through the sorted list, a sample is selected only if it hasn’t been excluded in the previous steps. The user can adjust key parameters, such as the reference sample, the initial radius *r*, the parameter *k*, and the distance metric parameter *p*. The computational cost of OptiSim and DISE is ∼𝒪(*nd*), making them more computationally tractable than the MaxMin and MaxSum methods.

#### 2.2.2 Partition-Based Selection Algorithms

Partition-based selection methods rely on partitioning the data space into regions (commonly portions of multi-dimensional hyper-cubes) based on various criteria, and (randomly) selecting samples from each region. While partition-based algorithms are suitable for a wider range of diversity-related tasks, such as identifying and filling in diversity voids and comparing diversities of different populations, they are only computationally practical when the feature space is low-dimensional.^14^

The “Equisized Independent” method divides the feature space into equal-sized bins along each dimension. The partitioning is done independently for each dimension, so the entire feature space is divided into rectangular or cubic segments where each segment has the same dimensional length. Similarly, the “Equisized Dependent” method also partitions the space into bins of equal size, but here, the size and position of bins in one dimension can depend on the partitioning of previous dimensions. This means that the order of features affects the outcome, as each dimension’s partitioning might adjust based on the dimensions that were processed before it. The “Equifrequent Independent” divides the feature space into bins, each containing approximately the same number of sample points. Unlike equisized approaches, the sizes of these bins may vary to maintain an equal frequency of points per bin. This partitioning occurs independently across each dimension, ensuring that each bin within a single dimension contains a similar number of points, irrespective of the bin configurations in other dimensions. In contrast, the “Equifrequent Dependent” method achieves equifrequent partitioning by considering dependency between dimensions. This interdependency ensures that each bin maintains an approximately equal distribution of sample points while taking into account the partitioning configurations of earlier dimensions.

#### 2.2.3 Similarity-Based Selection Methods

Similarity-based algorithms aim to minimize the average similarity of the selected sample, so they are closely related to the distance-based methods described in Section 2.2.1. These algorithms commonly use pairwise similarity measures, where the average similarity is calculated over all possible sample pairs. This step incurs a computational cost of *O*(*n*^2^) where *n* is the number of selected samples. This limits the feasible subset size that can be considered. To address this limitation, we implement the recently proposed extended similarity indices (eSIM)^70,71^ and instant similarity index (iSIM)^72^ families of manywise similarity measures. eSIM generalizes pairwise similarity metrics to the n-similarity setting, and iSIM compares multiple samples simultaneously and produces results nearly equivalent to average pairwise (dis)similarity, but at a significantly lower computational cost.^70,71^ Both methods start from a user-defined point and iteratively add the unselected element that results in the lowest average similarity to the current set. This continues until the target subset size is reached.

Given a feature matrix **X**_*s ×d*_, the eSIM and iSIM methods start by computing the columnwise sum. eSIM classifies (counts) each column comparing its sum of value with a threshold as **1-similarity, 0-similarity** or **dissimilarity** column, and then a weighting function is used to consider partial similarity and dissimilarity. The similarity metrics are then defined as functions of these counters. iSIM, on the other hand, computes the number of coinciding **on** or **off** pair of bits, and the number of mismatches in each column from the column-wise sum. The average similarity indices are then computed from these counters. The following similarity indices are supported: Austin-Colwell, Baroni-Urbani-Buser, Consoni-Todschini, Faith, Gleason, Jaccard, Jaccard 0-variant, Jaccard-Tanimoto, Rogers-Tanimoto, Russel-Rao, Sokal-Michener and Sokal-Sneath.^70^ These methods were initially designed to compare binary feature vectors, nonetheless, extending eSIM to continuous variables is straightforward.^73^ On the other hand, the mathematical formulation of the iSIM approach relies on binary feature representations, and therefore, it does not apply to non-binary features. However, both methods have 𝒪(*nd*) cost, and select a diverse set closely similar to the `MaxSum` method.

### 2.3 Similarity and Diversity Measures

Similarity measures quantify how alike two objects are, typically ranging from 0 (no similarity) to 1 (identical). Supporting functions are implemented to compute pairwise similarity matrices using measures such as the Jaccard-Tanimoto coefficient^74^ and its modified form for binary fingerprints.^75^ On the other hand, dissimilarity measures indicate how different two objects are, with lower values indicating greater similarity. Distances are a specific class of dissimilarity measures that satisfy non-negativity, symmetry, and the triangle inequality. `Selector` is fully compatible with the extensive distance functions available in `SciPy`^76^ and `Scikit-Learn`,^77^ and also includes functions for converting similarity scores into distance measures.^78^ These capabilities complement external tools for computing distance and similarity matrices.^78^

Diversity measures quantify the variability within a set and are categorized into two types: (1) absolute diversity measures, which depend solely on the feature matrix of the selected samples; and (2) relative diversity measures, which require both the full feature matrix and the indices of the selected subset, for example, hypersphere overlap.^5^ In the first category, `Selector` currently implements the log determinant function^42^, Shannon entropy^50,79^, explicit diversity index^80^, Gini coefficient^81^, and Wasserstein Distance to Uniform Distribution (WDUD)^42^. Among these, the hypersphere overlap, log-determinant, and WDUD methods support both binary and non-binary feature matrices, while the remaining metrics are restricted to binary features. Higher values of the log-determinant function, Shannon entropy, and explicit diversity index indicate greater diversity, whereas lower values of WDUD, hyper-sphere overlapping, and Gini coefficient are associated with increased diversity.

## 3 Code Examples

The code snippets below demonstrate how easily different selection methods generate diverse subsets from one or more clusters using either features or pairwise distances. Extended versions of these examples are provided as quick-start Jupyter notebooks in the repository; however, some of the output images are included in Figure SI 1 and Figure SI 2. For easy visualization, we use a 2D dataset generated from two isotropic Gaussian distributions of `Scikit-Learn`. Given a feature matrix **X**_*s×d*_ representing *s* samples in *d* dimensions or features, the corresponding pairwise distance matrix is denoted by **D**_*s× s*_. The `labels` argument indicates which of the two classes each data point belongs to.

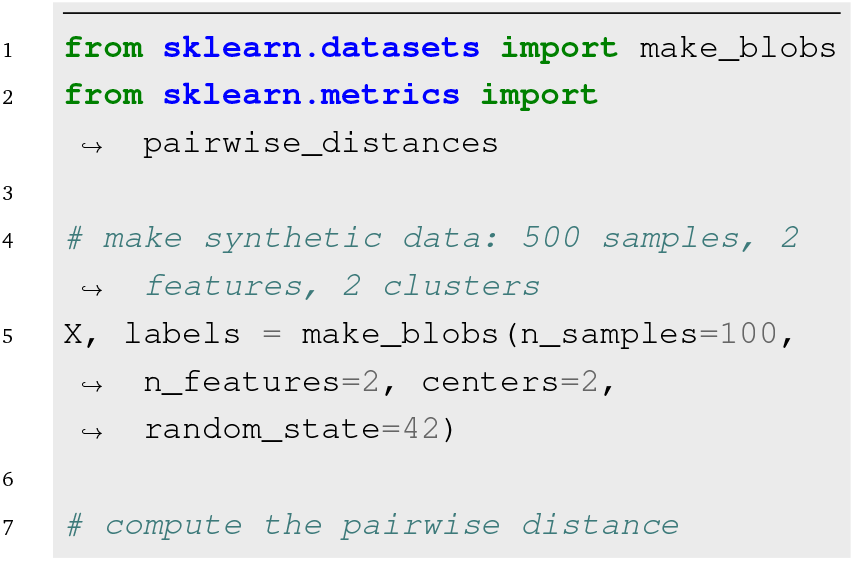
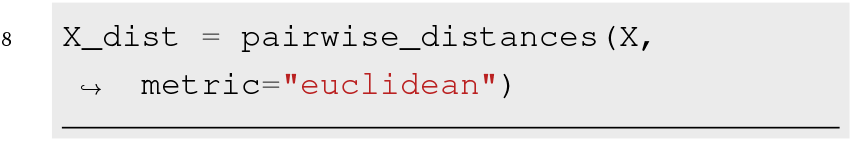

### Distance-Based Selection

Given a symmetric distance matrix, the `select` method of any of the distance-based classes can be used to choose a subset of the data with size *n*. For example,

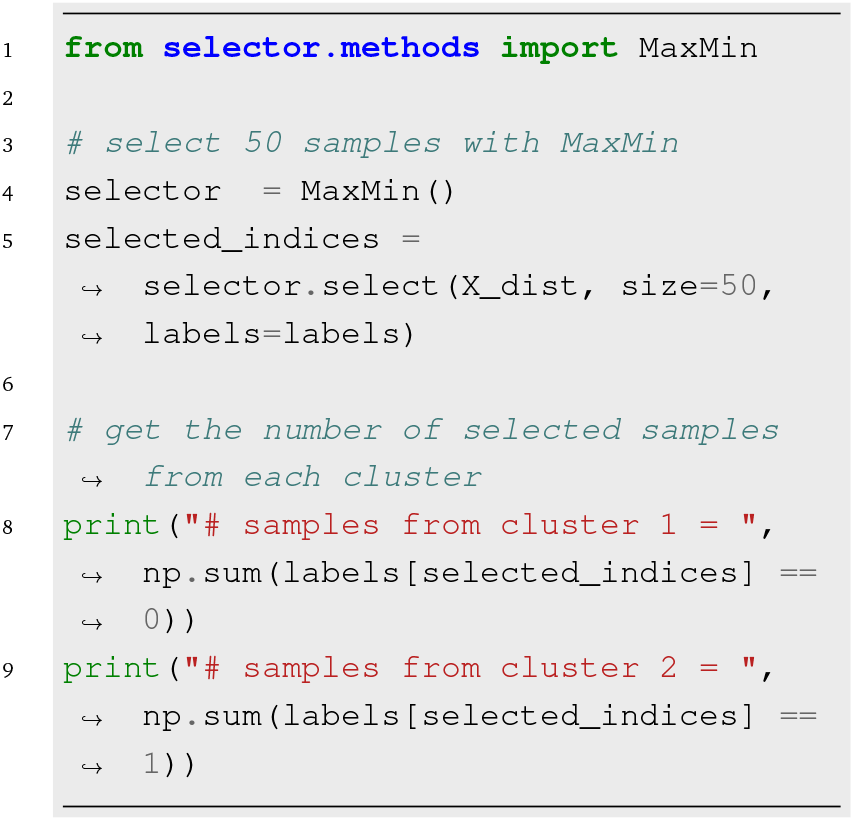

Providing the optional `label` argument guides the algorithm to select *n* data points while preserving the class distribution. Alternatively, a user-defined function `fun_dist` can be provided when initializing distance-based classes, in which case the input array of `select` method is treated as a feature matrix.

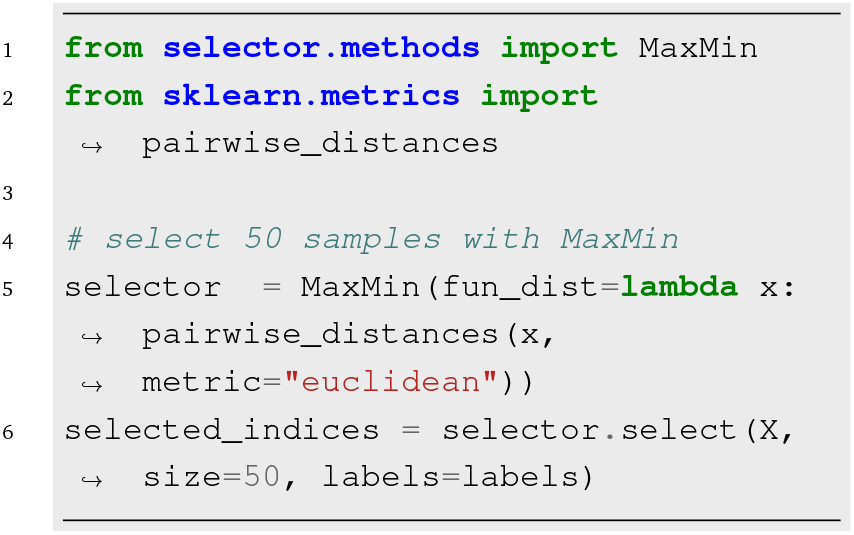

### Partition-Based Selection

The feature matrix **X**, which defines the space traversed by the data, is partitioned based on the specified `bin_method` argument, and in a similar fashion, the `select` method is used to choose a subset.

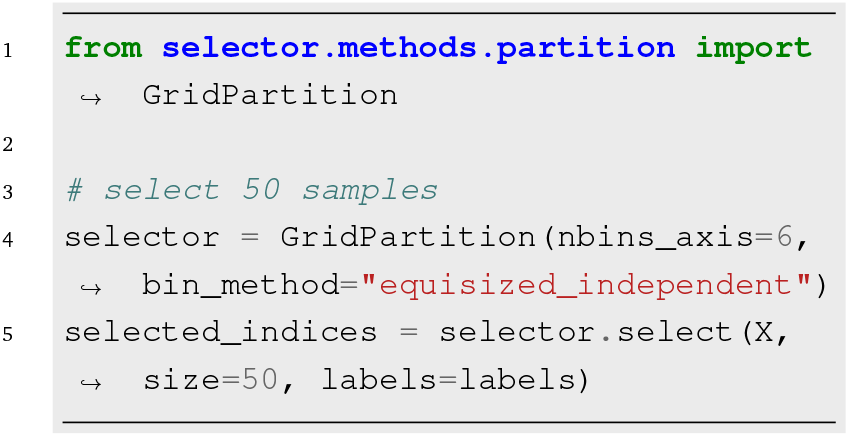

### Similarity-Based Selection

Currently, the `NSimilarity` supports subset selection using 16 implemented similarity measures. While originally designed for binary strings, support for non-binary data is available through the `eSIM` method by `preprocess_data=True`. For example,

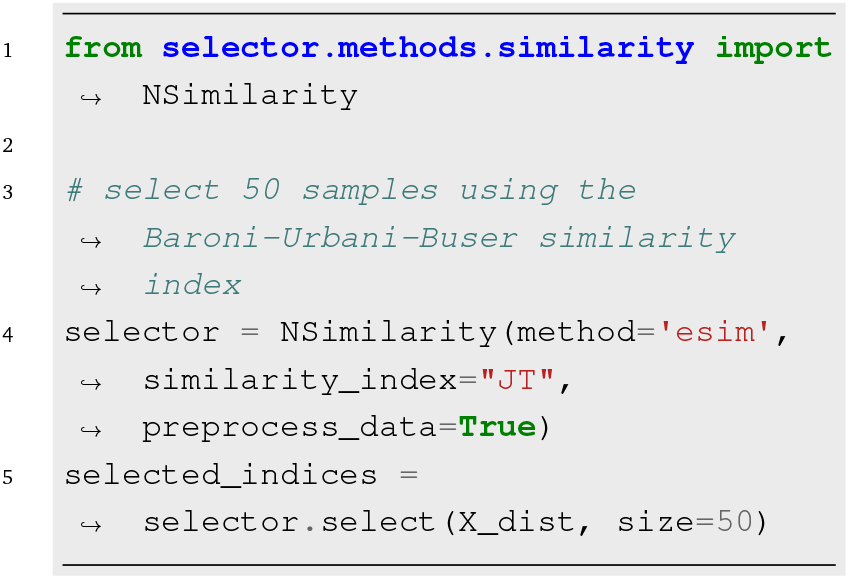

Figure SI 2 illustrates the selected points using the Jaccard-Tanimoto (JT) coefficient alongside three other similarity-based descriptors. All methods select the same set of points, which tend to lie on the periphery of the distribution. This is an expected outcome for average-based selection methods in lowdimensional spaces. However, the descriptors select different subsets in higher-dimensional feature spaces.

### Diversity Measures

The `diversity` module provides functions to compute diversity metrics given the full feature matrix and a selected subset.

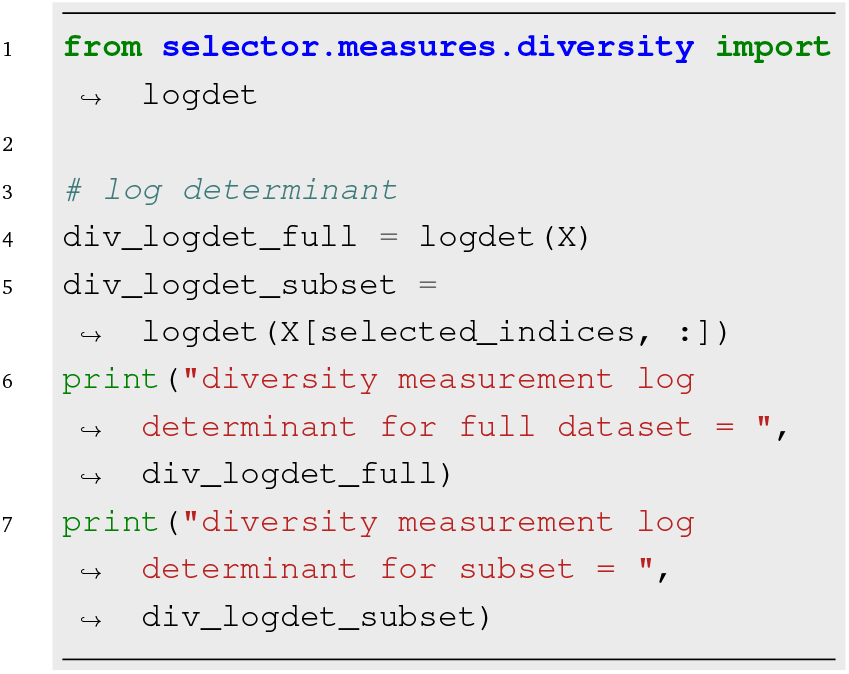

## 4 Applications

To demonstrate the versatility of the `Selector` library, we present several examples to showcase its flexibility and broad applicability across disciplines.

### 4.1 Selecting Representative Compounds from Biased Datasets

Biased datasets often overrepresent certain categories, making random sampling prone to reinforcing existing imbalances rather than capturing true diversity. We demonstrate how `Selector` can be used to extract a diverse subset from such biased datasets, for example, synthesized compounds or biological assays, which are costly and time-consuming to generate. To demonstrate this, we manually curated a biased dataset of 145 organic compounds, comprising 98 alkanes, 19 alcohols, 10 aldehydes, 9 ketones, 7 benzene derivatives, and 2 cycloalkanes (see Table SI 1), each represented with an Extended Connectivity Fingerprints (ECFP4). We then selected 9 compounds using MaxMin with the modified Tanimoto distance^75^ measure, without incorporating any information about compound classes. When selecting a subset of 9 compounds, Figure 2 shows that MaxMin successfully includes at least one compound from each class. In contrast, random sampling disproportionately favors the majority classes (alkanes and alcohols), overlooking aldehydes, ketones, benzene derivatives, and cycloalkanes. As expected, random selection becomes more likely to include underrepresented groups as the subset size increases. In short, a proper selection algorithm mitigates bias effects and ensures diversity across underrepresented classes.

**Figure 2.**
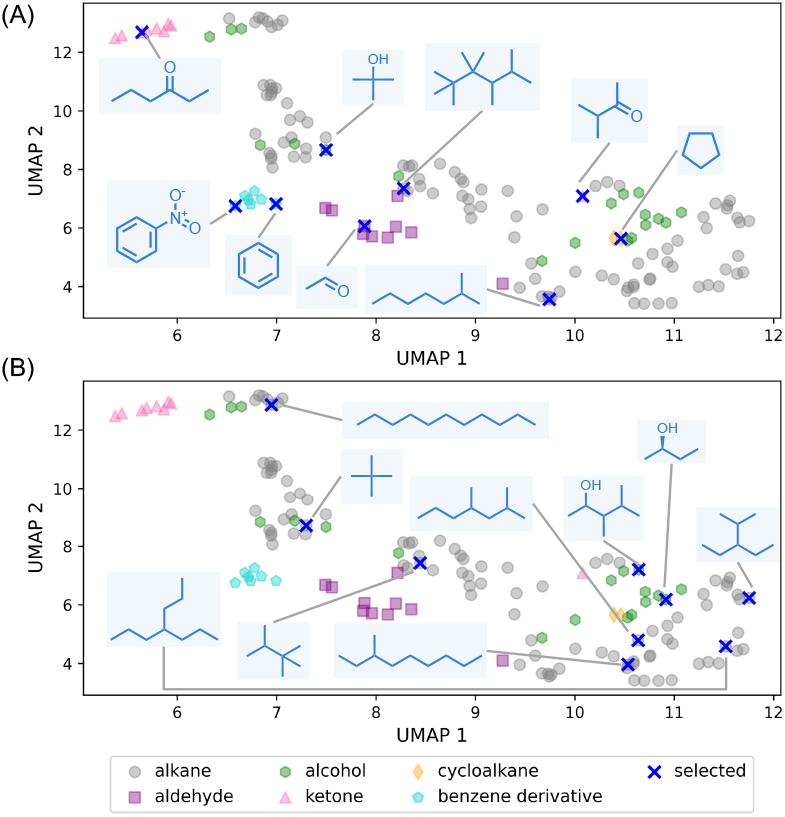
Selecting representative compounds from a biased dataset of 145 organic compounds (c.f. Table SI 1), each encoded using Extended Connectivity Fingerprints (ECFP4) and projected into two dimensions via Uniform Manifold Approximation and Projection (UMAP) for visualization. Blue crosses indicate the 9 selected samples in 512 dimensions using (A) the MaxMin algorithm and (B) random sampling. The ECFP4 fingerprint was calculated with `RDKit` ^48^ version 2024.03.5.

### 4.2 Sampling Potential Energy Surface

Machine learning–based modeling of potential energy surfaces (PES) has emerged as a powerful approach for accurate prediction of the energy landscapes of molecular systems.^82–87^ Training these models typically involves generating datasets by systematically or randomly sampling a high-dimensional PES of molecules, followed by quantum chemistry calculations.^88–90^ The resulting data are then divided into training, validation, and test subsets using either random or stratified sampling. To illustrate the significance of diverse subset selection, we constructed a four-well potential as described by^91,92^ (detailed in Section SI 1) in 2-dimensions, which allows us to easily visualize both the dataset and the selected subsets. To numerically represent the *V* (*q*_1_, *q*_2_) potential, we uniformly sampled 10 million (*q*_1_, *q*_2_) coordinates in two dimensions over the range [− 3.0, 3.0], which includes all four minima. Then, we used the Monte-Carlo (MC) to select 10,000 points from the Boltzmann distribution, exp(−1.5*k* × (*V* (*q*_1_, *q*_2_) − *V*_*D*_)), where *V*_*D*_ is the potential of the deepest minima. This generated two datasets of equal size but varying bias: the *k* = 10 dataset is strongly biased toward areas around the local minima (and entirely misses the shallow minimum, whereas the *k* = 2 dataset provides a more thorough sampling of the PES. The two datasets are marked with black stars in Figure 3. We applied various selection methods to select 1% and 64% of each dataset, representing small and large subsets, respectively.

**Figure 3.**
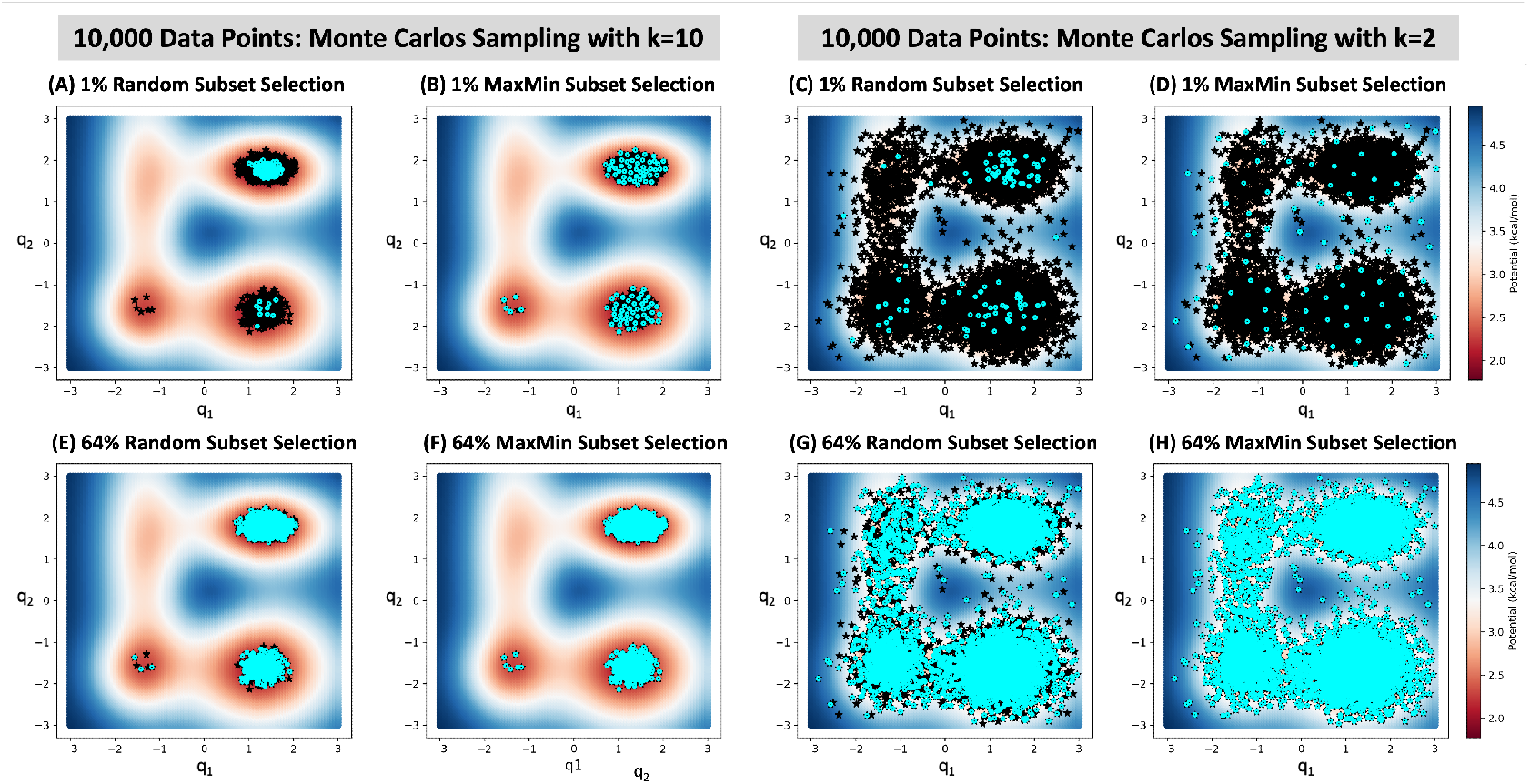
Selecting subsets (colored with cyan circles) of the four-well potential with Random (1st and 3rd columns) and MaxMin (2nd and 4th columns) methods from a dataset of 10,000 points (colored with black stars) generated with Monte Carlo sampling using values of k=10 (two left columns; strong bias) and k=2 (two right columns; weak bias). Similar plots showing subset selection with MaxSum, OptiSim, and Grid Partition methods are provided in Figure SI 5, Figure SI 6, and Figure SI 7.

Figure 3 (A) and (C) plots show that random sampling fails to yield a representative subset when selecting a small fraction of datasets, especially for a biased dataset. They reveal that random sampling overlooks less populated areas that are farthest from the local minima. Using stratified random selection improves the subsets, but it does not completely fix the issue. On the other hand, Figure 3 (B) and (D) show that the MaxMin algorithm consistently selects a balanced and diverse subset for both subset sizes. This contrast between random sampling and the MaxMin algorithm is less pronounced when a large subset of the data is selected, as shown in plots (E) to (H) in Figure 3. The results from MaxSum, OptiSim, and grid partition methods are presented in Figure SI 5, Figure SI 6, and Figure SI 7.As previously observed, MaxSum tends to select data points from peripheral regions, while partition-based methods often miss certain areas of the PES. In contrast, OptiSim selects more diverse subsets.

### 4.3 Selecting Molecules from a Chemical Database

Selecting diverse subsets of data is a crucial step in cheminformatics and data-driven modeling. To demonstrate `Selector` in this context, we applied various selection methods to extract 10–50% subsets from 27 ChEMBL datasets, which are well-curated libraries of biologically relevant molecules targeting diverse proteins^93^. The Figure 4 shows the performance of the ChEMBL204 Ki library, which includes 2754 compounds with inhibitory activity against thrombin. Based on the log-determinant diversity measure, MaxMin and OptiSim are the best-performing selection methods, while GridPartition is the least effective.

**Figure 4.**
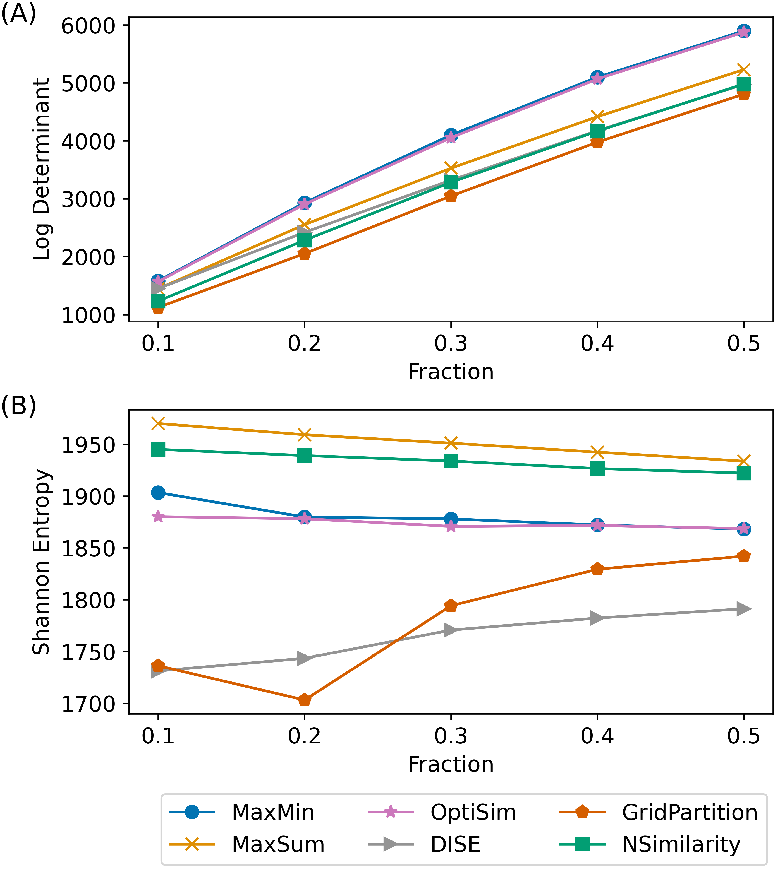
Diversity measured by (A) Log-determinant and (B) Shannon entropy for selected fractions of the ChEMBL204 Ki dataset (composed of 2754 compounds) using various selection methods. The 2048-bit RDKit fingerprint was generated with `Chem.RDKFingerprint` in `RDKit` ^48^.

In contrast, Shannon entropy shows that MaxSum and NSimilarity consistently result in the highest diversity across all subset fractions. Grid partitioning performs worst when less than 30% of the data is selected, while DISE shows the lowest diversity for larger fractions. Figure SI 8 presents similar plots for the WDUD and Gini coefficient measures. Our results show that NSimilarity and MaxSum tend to guarantee a larger “volume” of the selected subsets^71^. The discrepancy between log-determinant and other diversity measures suggests that one should choose proper diversity measures based on the problem at hand, as there is no universally accepted diversity measure.

GridPartition tends to underperform, largely due to the high dimensionality of the molecular representation (2048 bits), making it more suitable for low-dimensional data^14^. The DISE algorithm is sensitive to the choice of initial reference molecule and radius^69^, which can result in low-diversity selections. Further investigation into optimal radius settings may help improve its performance. Among the diversity metrics evaluated, the log-determinant shows a notably different trend; prior work has highlighted its limitations in capturing diversity^42,94^. As the selected subset fraction increases, diversity scores of selection methods begin to converge, reflecting the diversity of the full dataset. This trend holds consistently across all ChEMBL subsets. Additional results for the remaining datasets are discussed in the Supporting Information, see Figure SI 9–SI 34.

## 5 Conclusions

This paper presents `Selector`, a free and open-source Python package for selecting diverse subsets using distance-based, partition-based, and similarity-based algorithms as well as computing diversity measures. It integrates seamlessly with machine learning and data analysis libraries, offering users extensive flexibility to customize selection strategies to fit their specific needs. It is designed to be easy for users to apply to their problems and straightforward for developers to extend with new algorithms. In addition to interactive tutorial notebooks, we have developed a web interface that enables users to apply `Selector` to their data by simply uploading it, making the tool accessible without requiring coding or local installation.

Even though the examples presented focus on applications in computational chemistry and cheminformatics, the challenge of representative subset selection is broadly relevant across scientific and engineering domains, especially with the rise of data-driven modeling. Diversity-aware sampling remains underutilized in many studies, which motivated the development of `Selector` as a flexible and user-friendly toolkit for customizable diversity-based sampling. It is developed following modern software development practices to ensure both accessibility and longterm sustainability. The QC-Devs team remains committed to its ongoing maintenance and development, and we welcome feedback and encourage contributions from the community.

## Data and Software Availability

The source code is available at https://github.com/theochem/Selector, and the documentation (including tutorial notebooks) can be accessed at https://selector.qcdevs.org/. For codes and data to reproduce the figures, please refer to https://figshare.com/articles/figure/data_for_reproducing_figures/26886652^95^. The web interface of the `Selector` is available at https://huggingface.co/spaces/QCDevs/Selector.

## Author Contributions

Fanwang Meng: Conceptualization, Software, Validation, Writing – original draft, review & editing; Marco Martínez González: Software, Validation, Writing – original draft, review & editing Valerii Chuiko: Software, Validation, Logo Design; Alireza Tehrani: Software, Validation; Abdul Rahman Al Nabulsi: Software; Abigail Broscius: Software; Hasan Khaleel: Software; Kenneth López-Pérez: Writing – original draft; Ramón Alain Miranda-Quintana: Software, Writing – original draft, review & editing, Funding acquisition; Paul W. Ayers: Conceptualization, Software, Validation, Writing – review & editing, and Funding acquisition; Farnaz Heidar-Zadeh: Conceptualization, Software, Validation, Writing – original draft, review & editing, and Funding acquisition.

## Acknowledgements

F.H. acknowledges financial support from the Natural Sciences and Engineering Research Council (NSERC) of Canada, the Digital Research Alliance of Canada, and the Center for Advanced Computing (CAC) at Queen’s University. P.W.A. acknowledges NSERC, the Canada Research Chairs, and the Digital Research Alliance of Canada for financial and computational support. R.A.M.Q. and K.L.P. thank the National Institute of General Medical Sciences of the National Institutes of Health under award number R35GM150620. F.M. acknowledges the Banting Postdoctoral Fellowship administered by Canada’s three research granting agencies: the Canadian Institutes of Health Research (CIHR), the Natural Sciences and Engineering Research Council of Canada (NSERC) and the Social Sciences and Humanities Research Council (SSHRC). F.M. also thanks the support of the Alliance’s DRI (Digital Research Infrastructure) EDIA (Equity, Diversity, Inclusion and Accessibility) Champions Pilot Program, Digital Research Alliance of Canada, for financial and computational support. The authors also acknowledge the helpful discussions and help with this package with Maximilian van Zyl (Queen’s University), and Yang Xu (McMaster University).

## Supporting Information Available

Supporting information is available free of charge at https://pubs.acs.org/doi/10.1021/acs.jcim/xxx.

## TOC Graphic

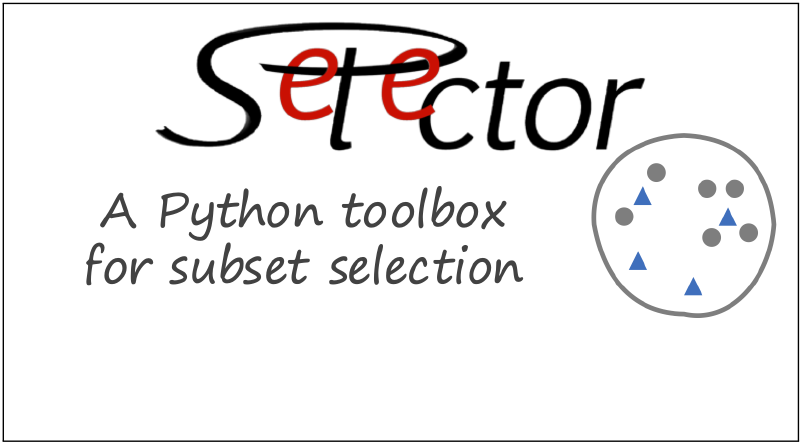

