## Supporting Information for "Selector: A General Python Library for Diverse Subset Selection"

### List of Tables

SI 1 SMILES strings for the manually-curated biased chemical library with 145 compounds, which contains 98 alkanes, 19 alcohols, 10 aldehydes, 9 ketones, 7 benzene derivatives and 2 cycloalkanes. . . . . . . . . . . . . . . . . 10

### List of Figures

SI 1 Comparison of distance-based (A) and partition-based (B) selectors for selecting 50 samples from a set of size 500 in 2D dimensions. For (B), The equis-(in)dependent and equif-(in)dependent labels refer to the (in)dependent variants of the equisize and equifrequent grid partitioning method. (A) shows that MaxSum algorithm selects samples from the peripheral space occupied by the data points. When the number of selected samples $s$ is fewer than the number of dimensions $d$, this ensures that each dimension is adequately represented in the chosen subset. . . . . . . . . . . . . . . . . . . 9

SI 2 Comparing NSimilarity based selection methods. . . . . . . . . . . . . . . . . 17

SI 3 Running time comparisons of different subset selection algorithms. The 1024-bit ECFP4 fingerprints were computed with `RDKit` for the 14k logP dataset from EPA public data[1]. The `MaxMinPicker` method in `RDKit` was as the base line. It is noticed OptiSim are slower because the setup of searching the optimal radius in `Selector` in order to get the required number of selected data points. . . . . . . . . . . . . . . . . . . . . . . . . . . . . . 17

SI 4 The four-well potential with four minima: $V_A(-1.17, 1.56) = 2.84$, $V_B(-1.29, -1.53) = 2.28$, $V_C(1.29, -1.65) = 2.01$, and $V_D(1.4, 1.78) = 1.77$ kcal/mol.[2] . . . . . . . 18

SI 5 Selecting subsets (colored with cyan circles) of four-well potential with MaxSum (1st and 3rd columns) and OptiSim (2nd and 4th columns) methods from a dataset of 10,000 points (colored with black stars) generated with Monte Carlo sampling using values of k=10 (two left columns) and k=2 (two right columns). . . . . . . . . . . . . . . . . . . . . . . . . . . . . 18
SI 6 Selecting subsets (colored with cyan circles) of four-well potential with Grid Partition Equifrequent Dependent (1st and 3rd columns) and Independent (2nd and 4th columns) methods from a dataset of 10,000 points (colored with black stars) generated with Monte Carlo sampling using values of k=10 (two left columns) and k=2 (two right columns). For the Grid Partition, 5 bins are used along each access. . . . . . . . . . . . . . . . . . . . 19
SI 7 Selecting subsets (colored with cyan circles) of four-well potential with Grid Partition Equisize Dependent (1st and 3rd columns) and Independent (2nd and 4th columns) methods from a dataset of 10,000 points (colored with black stars) generated with Monte Carlo sampling using values of k=10 (two left columns) and k=2 (two right columns). For the Grid Partition, 5 bins are used along each access. . . . . . . . . . . . . . . . . . . . . . . . . . . . 19
SI 8 Diversity selection measurements of ChEMBL204 Ki subset ($n_{total} = 2754$) selection with different selection methods. (A) Log-determinant of the selected subset. (B) Wasserstein distance to uniform distribution (WDUD) of the selected subset. (C) Shannon entropy of the selected subset. (D) Gini coefficient of the selected subset. Based on the WDUD, MaxSum and NSimilarity perform the best across the different fractions. Grid partitioning has the lowest diversity when less than 30% of the data is selected, but for larger fractions of the data DISE performs worst. The Gini coefficient, MaxSum and NSimilarity consistently yield the most diverse sets and DISE consistently gives the least diverse set. . . . . . . . . . . . . . . . . . . . . . . . 20

SI 9 Diversity selection measurements of ChEMBL1871 Ki subset ($n_{total} = 659$) selection with different selection methods. (A) Log-determinant of the selected subset. (B) Wasserstein distance to uniform distribution (WDUD) of the selected subset. (C) Shannon entropy of the selected subset. (D) Gini coefficient of the selected subset. . . . . . . . . . . . . . . . . . . . . . . . . 21
SI 10 Diversity selection measurements of ChEMBL2034 Ki subset ($n_{total} = 750$) selection with different selection methods. (A) Log-determinant of the selected subset. (B) Wasserstein distance to uniform distribution (WDUD) of the selected subset. (C) Shannon entropy of the selected subset. (D) Gini coefficient of the selected subset. . . . . . . . . . . . . . . . . . . . . . . . . 22
SI 11 Diversity selection measurements of ChEMBL2047 EC50 subset ($n_{total} = 631$) selection with different selection methods. (A) Log-determinant of the selected subset. (B) Wasserstein distance to uniform distribution (WDUD) of the selected subset. (C) Shannon entropy of the selected subset. (D) Gini coefficient of the selected subset. . . . . . . . . . . . . . . . . . . . . . . . . 23
SI 12 Diversity selection measurements of ChEMBL2147 Ki subset ($n_{total} = 1456$) selection with different selection methods. (A) Log-determinant of the selected subset. (B) Wasserstein distance to uniform distribution (WDUD) of the selected subset. (C) Shannon entropy of the selected subset. (D) Gini coefficient of the selected subset. . . . . . . . . . . . . . . . . . . . . . . . . 24
SI 13 Diversity selection measurements of ChEMBL214 Ki subset ($n_{total} = 3317$) selection with different selection methods. (A) Log-determinant of the selected subset. (B) Wasserstein distance to uniform distribution (WDUD) of the selected subset. (C) Shannon entropy of the selected subset. (D) Gini coefficient of the selected subset. . . . . . . . . . . . . . . . . . . . . . . . . 25

SI 14 Diversity selection measurements of ChEMBL218 EC50 subset ($n_{total} = 1031$) selection with different selection methods. (A) Log-determinant of the selected subset. (B) Wasserstein distance to uniform distribution (WDUD) of the selected subset. (C) Shannon entropy of the selected subset. (D) Gini coefficient of the selected subset. . . . . . . . . . . . . . . . . . . . . . . . . . 26
SI 15 Diversity selection measurements of ChEMBL219 Ki subset ($n_{total} = 1859$) selection with different selection methods. (A) Log-determinant of the selected subset. (B) Wasserstein distance to uniform distribution (WDUD) of the selected subset. (C) Shannon entropy of the selected subset. (D) Gini coefficient of the selected subset. . . . . . . . . . . . . . . . . . . . . . . . . . 27
SI 16 Diversity selection measurements of ChEMBL228 Ki subset ($n_{total} = 1704$) selection with different selection methods. . . . . . . . . . . . . . . . . . . . . 28
SI 17 Diversity selection measurements of ChEMBL231 Ki subset ($n_{total} = 973$) selection with different selection methods. (A) Log-determinant of the selected subset. (B) Wasserstein distance to uniform distribution (WDUD) of the selected subset. (C) Shannon entropy of the selected subset. (D) Gini coefficient of the selected subset. . . . . . . . . . . . . . . . . . . . . . . . . . 29
SI 18 Diversity selection measurements of ChEMBL233 Ki subset ($n_{total} = 3142$) selection with different selection methods. (A) Log-determinant of the selected subset. (B) Wasserstein distance to uniform distribution (WDUD) of the selected subset. (C) Shannon entropy of the selected subset. (D) Gini coefficient of the selected subset. . . . . . . . . . . . . . . . . . . . . . . . . . 30
SI 19 Diversity selection measurements of ChEMBL234 Ki subset ($n_{total} = 3657$) selection with different selection methods. (A) Log-determinant of the selected subset. (B) Wasserstein distance to uniform distribution (WDUD) of the selected subset. (C) Shannon entropy of the selected subset. (D) Gini coefficient of the selected subset. . . . . . . . . . . . . . . . . . . . . . . . . . 31

SI 20 Diversity selection measurements of ChEMBL235 EC50 subset ($n_{total} = 2349$) selection with different selection methods. (A) Log-determinant of the selected subset. (B) Wasserstein distance to uniform distribution (WDUD) of the selected subset. (C) Shannon entropy of the selected subset. (D) Gini coefficient of the selected subset. . . . . . . . . . . . . . . . . . . . . . . . . . 32
SI 21 Diversity selection measurements of ChEMBL236 Ki subset ($n_{total} = 2598$) selection with different selection methods. (A) Log-determinant of the selected subset. (B) Wasserstein distance to uniform distribution (WDUD) of the selected subset. (C) Shannon entropy of the selected subset. (D) Gini coefficient of the selected subset. . . . . . . . . . . . . . . . . . . . . . . . . . 33
SI 22 Diversity selection measurements of ChEMBL237 EC50 subset ($n_{total} = 955$) selection with different selection methods. . . . . . . . . . . . . . . . . . . . . . 34
SI 23 Diversity selection measurements of ChEMBL237 Ki subset ($n_{total} = 2602$) selection with different selection methods. (A) Log-determinant of the selected subset. (B) Wasserstein distance to uniform distribution (WDUD) of the selected subset. (C) Shannon entropy of the selected subset. (D) Gini coefficient of the selected subset. . . . . . . . . . . . . . . . . . . . . . . . . . 35
SI 24 Diversity selection measurements of ChEMBL238 Ki subset ($n_{total} = 1052$) selection with different selection methods. (A) Log-determinant of the selected subset. (B) Wasserstein distance to uniform distribution (WDUD) of the selected subset. (C) Shannon entropy of the selected subset. (D) Gini coefficient of the selected subset. . . . . . . . . . . . . . . . . . . . . . . . . . 36
SI 25 Diversity selection measurements of ChEMBL239 EC50 subset ($n_{total} = 1721$) selection with different selection methods. (A) Log-determinant of the selected subset. (B) Wasserstein distance to uniform distribution (WDUD) of the selected subset. (C) Shannon entropy of the selected subset. (D) Gini coefficient of the selected subset. . . . . . . . . . . . . . . . . . . . . . . . . . 37

SI 26 Diversity selection measurements of ChEMBL244 Ki subset ($n_{total} = 3097$) selection with different selection methods. (A) Log-determinant of the selected subset. (B) Wasserstein distance to uniform distribution (WDUD) of the selected subset. (C) Shannon entropy of the selected subset. (D) Gini coefficient of the selected subset. . . . . . . . . . . . . . . . . . . . . . . . . 38
SI 27 Diversity selection measurements of ChEMBL264 Ki subset ($n_{total} = 2862$) selection with different selection methods. (A) Log-determinant of the selected subset. (B) Wasserstein distance to uniform distribution (WDUD) of the selected subset. (C) Shannon entropy of the selected subset. (D) Gini coefficient of the selected subset. . . . . . . . . . . . . . . . . . . . . . . . . 39
SI 28 Diversity selection measurements of ChEMBL2835 Ki subset ($n_{total} = 615$) selection with different selection methods. (A) Log-determinant of the selected subset. (B) Wasserstein distance to uniform distribution (WDUD) of the selected subset. (C) Shannon entropy of the selected subset. (D) Gini coefficient of the selected subset. . . . . . . . . . . . . . . . . . . . . . . . . 40
SI 29 Diversity selection measurements of ChEMBL287 Ki subset ($n_{total} = 1328$) selection with different selection methods. (A) Log-determinant of the selected subset. (B) Wasserstein distance to uniform distribution (WDUD) of the selected subset. (C) Shannon entropy of the selected subset. (D) Gini coefficient of the selected subset. . . . . . . . . . . . . . . . . . . . . . . . . 41
SI 30 Diversity selection measurements of ChEMBL2971 Ki subset ($n_{total} = 976$) selection with different selection methods. (A) Log-determinant of the selected subset. (B) Wasserstein distance to uniform distribution (WDUD) of the selected subset. (C) Shannon entropy of the selected subset. (D) Gini coefficient of the selected subset. . . . . . . . . . . . . . . . . . . . . . . . . 42

SI 31 Diversity selection measurements of ChEMBL3979 EC50 subset ($n_{total}$ = 1125) selection with different selection methods. (A) Log-determinant of the selected subset. (B) Wasserstein distance to uniform distribution (WDUD) of the selected subset. (C) Shannon entropy of the selected subset. (D) Gini coefficient of the selected subset. . . . . . . . . . . . . . . . . . . . . . . . . 43
SI 32 Diversity selection measurements of ChEMBL4005 Ki subset ($n_{total}$ = 960) selection with different selection methods. (A) Log-determinant of the selected subset. (B) Wasserstein distance to uniform distribution (WDUD) of the selected subset. (C) Shannon entropy of the selected subset. (D) Gini coefficient of the selected subset. . . . . . . . . . . . . . . . . . . . . . . . . 44
SI 33 Diversity selection measurements of ChEMBL4616 EC50 subset ($n_{total}$ = 682) selection with different selection methods. (A) Log-determinant of the selected subset. (B) Wasserstein distance to uniform distribution (WDUD) of the selected subset. (C) Shannon entropy of the selected subset. (D) Gini coefficient of the selected subset. . . . . . . . . . . . . . . . . . . . . . . . . 45
SI 34 Diversity selection measurements of ChEMBL4792 Ki subset ($n_{total}$ = 1471) selection with different selection methods. (A) Log-determinant of the selected subset. (B) Wasserstein distance to uniform distribution (WDUD) of the selected subset. (C) Shannon entropy of the selected subset. (D) Gini coefficient of the selected subset. . . . . . . . . . . . . . . . . . . . . . . . . 46

### SI 1 Four-Well Potential Construction

We used the four-well potential, plotted in Figure SI 4, which is described by:[2,3]

$$V(q_1,q_2) = V_0 + \sum_{i=0}^{4} a_i \exp[-\sigma_i^{q_1}(q_1-\alpha_i)^2 - \sigma_i^{q_2}(q_2-\beta_i)^2] \quad (1)$$

where $V_0 = 5.0$ kcal/mol, vector $a = [0.6, 3.0, 1.5, 3.2, 2.0]^T$ in kcal/mol, vector $\sigma^{q_1} = [1.0, 0.3, 1.0, 0.4, 1.0]^T$ in Å$^{-2}$, vector $\sigma^{q_2} = [1.0, 0.4, 1.0, 1.0, 0.1]^T$ in Å$^{-2}$, vector $\alpha = [0.1, 1.3, -1.5, 1.4, -1.3]^T$ in Å, and vector $\beta = [0.1, -1.6, -1.7, 1.8, 1.23]^T$ in Å. The four local minima labeled in Figure SI 4 correspond to $V_A(-1.17, 1.56) = 2.8450$, $V_B(-1.29, -1.53) = 2.2779$, $V_C(1.29, -1.65) = 2.0082$, and $V_D(1.4, 1.78) = 1.7755$ kcal/mol.

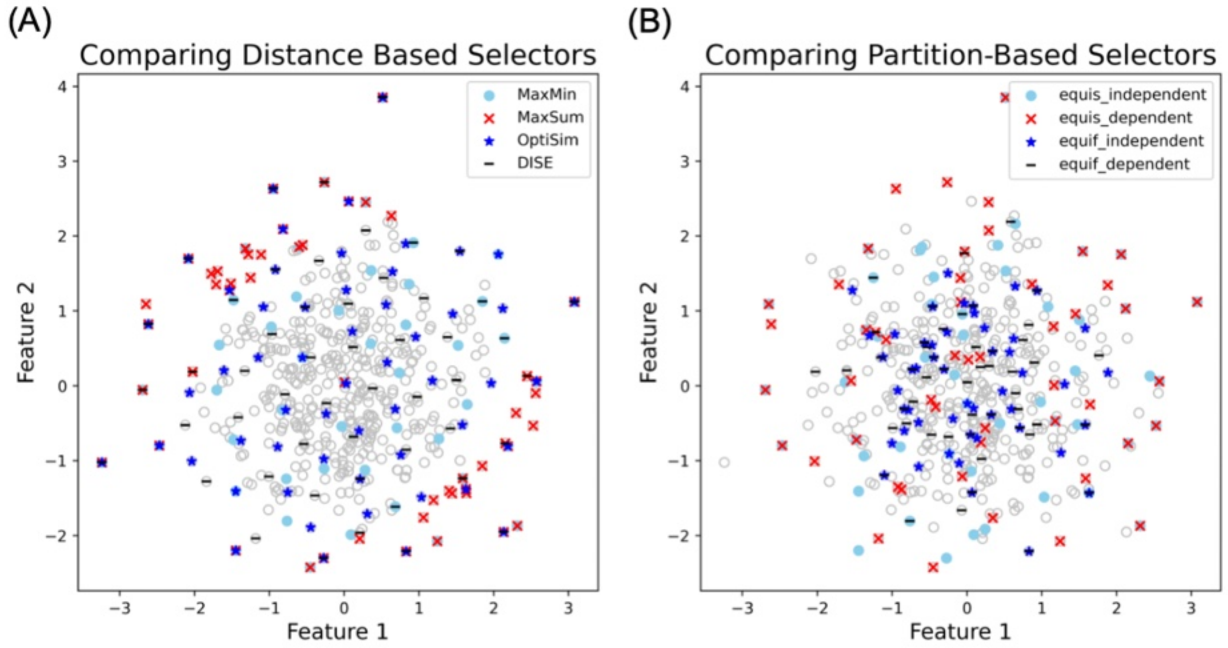


Fig. SI 1 Comparison of distance-based (A) and partition-based (B) selectors for selecting 50 samples from a set of size 500 in 2D dimensions. For (B), The equis-(in)dependent and equif-(in)dependent labels refer to the (in)dependent variants of the equisize and equifrequent grid partitioning method. (A) shows that MaxSum algorithm selects samples from the peripheral space occupied by the data points. When the number of selected samples $s$ is fewer than the number of dimensions $d$, this ensures that each dimension is adequately represented in the chosen subset.

Table SI 1 SMILES strings for the manually-curated biased chemical library with 145 compounds, which contains 98 alkanes, 19 alcohols, 10 aldehydes, 9 ketones, 7 benzene derivatives and 2 cycloalkanes.

| NO. | SMILES | Category |
|---|---|---|
| 1 | ketone | CCCCCC(=O)CC |
| 2 | aldehyde | CCC(C)C=O |
| 3 | benzene derivative | Nc1ccccc1 |
| 4 | ketone | CCCC(C)=O |
| 5 | alkane | CCC(CC)(C(C)C)C(C)(C)C |
| 6 | alkane | CCCCC(CCC)CCCC |
| 7 | alkane | CCCC(CC)CC(C)(C)C |
| 8 | alkane | CCCCCCCC(C)(C)C |
| 9 | alkane | CCC(C)CC(CC)CC(C)C |
| 10 | alkane | CCCC(C)CC(C)C |
| 11 | alkane | CCC(C)C(C)(C)CC |
| 12 | benzene derivative | Clc1ccccc1 |
| 13 | alkane | CCCCCC(C)C |
| 14 | alcohol | CCC(C)C(C)O |
| 15 | alkane | CCCC(CCC)CCC |
| 16 | alkane | CCCCCC |
| 17 | alkane | CCCCC(C)(C)CC(C)(C)C |
| 18 | cycloalkane | C1CCCCC1 |
| 19 | alkane | CC(C)(C)CCCCC(C)(C)C |
| 20 | alkane | CCCCCC(CC)CCCC |
| 21 | aldehyde | CCC=O |
| 22 | alkane | CCC(C)(C)CC |

Continued on next page

Table SI 1 – Continued from previous page

| NO. | SMILES | Category |
|---|---|---|
| 23 | alkane | CCCCC |
| 24 | alkane | CCC(CC)C(C)(C)C |
| 25 | alkane | CCCCCCC |
| 26 | alkane | CCC(CC)CC |
| 27 | alkane | CC(C)CC(C)C(C)(C)C |
| 28 | alkane | CC[C@@H](C)C(C)C |
| 29 | alkane | CCC(C)C |
| 30 | alkane | CCCCCC(C)CC |
| 31 | alkane | CC(C)C(C)(C)C(C)(C)C |
| 32 | alcohol | CC(C)C(C)O |
| 33 | alkane | CCCCCCCCC(C)(C)C |
| 34 | benzene derivative | NCc1ccccc1 |
| 35 | alcohol | CC(C)(C)O |
| 36 | alcohol | CCCCCO |
| 37 | alkane | CC(C)CC(C)(C)C |
| 38 | alcohol | CCC(O)CC |
| 39 | alcohol | CCCO |
| 40 | alkane | CCCCCCCCC |
| 41 | alcohol | CC[C@@H](C)CO |
| 42 | alcohol | CCC[C@@H](C)O |
| 43 | alcohol | CCCCO |
| 44 | alkane | CCCC(C)(C)C |
| 45 | alkane | CCCCC(C)CC |
| 46 | alkane | CCCCC[C@@H](C)CC(C)C |

Continued on next page

Table SI 1 – Continued from previous page

| NO. | SMILES | Category |
|---|---|---|
| 47 | alkane | CC(C)C(C(C)C)C(C)C |
| 48 | alkane | CCC(C)CCC(C)CC |
| 49 | alcohol | CC(C)CCO |
| 50 | ketone | CC(=O)C(C)C |
| 51 | benzene derivative | O=[N+]([O-])c1ccccc1 |
| 52 | aldehyde | CC(C)CC(C)C=O |
| 53 | alkane | CCC(C)C(C)(C)C(C)(C)C |
| 54 | alkane | CCCCC(C)C |
| 55 | alkane | CCCCCCCCC(C)C |
| 56 | alkane | CC(C)CCCCCC(C)C |
| 57 | alkane | CCCCCCCCCC(C)C |
| 58 | alkane | CCCCC(C)C(C)CC |
| 59 | alkane | CCC(C)CCCC(C)C |
| 60 | alkane | CCC(CC)C(C)C(CC)CC |
| 61 | alkane | CCC(C)C(C)(C)C |
| 62 | alcohol | CC(C)C(C)C(C)O |
| 63 | alkane | CCCC(C)CC |
| 64 | alkane | CCC(C)(C)C(C)C |
| 65 | alcohol | CC[C@H](C)CO |
| 66 | ketone | CCCCCCCC(=O)CC |
| 67 | alkane | CCC(CC)(CC)CC(C)C |
| 68 | alkane | CCCCCCC(C)(C)C |
| 69 | alkane | CCCCC(C)(C)C |
| 70 | alkane | CCCC(C)(C)CC |

Continued on next page

Table SI 1 – Continued from previous page

| NO. | SMILES | Category |
|---|---|---|
| 71 | alkane | CCCCCCC(C)C(C)CC |
| 72 | alkane | CCC(CC)C(C)C |
| 73 | alkane | CCCCCCCCCCC |
| 74 | alkane | CCCCCCCC(C)CC |
| 75 | ketone | CCC(C)=O |
| 76 | alkane | CC(C)C(C)C(C)(C)C |
| 77 | alcohol | CC(C)CO |
| 78 | benzene derivative | CCc1ccccc1 |
| 79 | alkane | CCC(CC)(CC)C(C)(C)C |
| 80 | alkane | CCCC(CC)CC |
| 81 | alkane | CCCC(CC)(CC)CC |
| 82 | benzene derivative | Cc1ccccc1 |
| 83 | alkane | CC[C@H](C)C(C)C |
| 84 | ketone | CCCCCC(=O)CCCC |
| 85 | alkane | CCCCC(C)C(C)C |
| 86 | alkane | CCCC(C)CCC |
| 87 | alkane | CCC(C)(C)CC(C)CC(C)C |
| 88 | benzene derivative | c1ccccc1 |
| 89 | aldehyde | CCCC=O |
| 90 | alkane | CCCCCCCCC(C)CC |
| 91 | alkane | CC(C)C(C)C(C)(C)C(C)(C)C |
| 92 | alkane | CCCCCCCC |
| 93 | alkane | CCCCCC(C)(C)C |
| 94 | alkane | CC(C(C)(C)C)C(C)(C)C |

Continued on next page

Table SI 1 – Continued from previous page

| NO. | SMILES | Category |
|---|---|---|
| 95 | alkane | CCC(CC)(CC)CC |
| 96 | alkane | CCC(C)(C)C |
| 97 | alkane | CC(C)(C)C(C)(C)C |
| 98 | alcohol | CC(C)(C)CO |
| 99 | alkane | CCC(C)(C)C(CC)(CC)CC |
| 100 | alcohol | CCC(C)(C)O |
| 101 | alkane | CC(C)CC(C)C |
| 102 | aldehyde | CC=O |
| 103 | alkane | CCC(C)(C)C(C)(C)CC |
| 104 | alkane | CC(C)C(C)C |
| 105 | alkane | CCC(C)C(C)(CC)C(C)(C)C |
| 106 | alkane | CC(CC(C)(C)C)C(C)(C)C |
| 107 | alkane | CCC(C)C(C)C(C)C(C)CC |
| 108 | aldehyde | CC(C)CC=O |
| 109 | alkane | CCCC(C)(CC)CC |
| 110 | alkane | CCCCCCCCCCC |
| 111 | alkane | CCCCC(C)C(C)CCCC |
| 112 | aldehyde | CC(C)C(C)C(C)(C)C=O |
| 113 | ketone | CCCC(=O)CC |
| 114 | alkane | CCCCCCC(C)CCCC |
| 115 | aldehyde | CC(C)C(C)C(C)C=O |
| 116 | alkane | CCC(C)CC |
| 117 | alkane | CC(C)CCCC(C)C |
| 118 | alcohol | CC(C)O |

Continued on next page

Table SI 1 – Continued from previous page

| NO. | SMILES | Category |
|---|---|---|
| 119 | alkane | CCC(C)(CC)C(C)(C)C |
| 120 | aldehyde | CC(C)CCCCC=O |
| 121 | alkane | CCC(CC)C(CC)CC |
| 122 | alkane | CC(C)(C)CC(C)(C)C |
| 123 | alkane | CCC(C)CC(C)(C)CC |
| 124 | alkane | CC(C)C(C)(C)C |
| 125 | alkane | CC(C)CCC(C)(C)C |
| 126 | alkane | CCCC(C)C |
| 127 | alkane | CC(C)CCC(C)C |
| 128 | alkane | CC(C)CC(C)(C)CC(C)C |
| 129 | alcohol | CC[C@@H](C)O |
| 130 | alkane | CC[C@@H](C)[C@@H](C)CC |
| 131 | aldehyde | CC(C)C(C)C=O |
| 132 | alkane | CCC(C)(C)C(C)(C)C |
| 133 | alkane | CC(C)C(C)C(C)C |
| 134 | alkane | CCCCCCC(C)C |
| 135 | alkane | CC(C)(C)C |
| 136 | alkane | CCCCC(CC)CCCC |
| 137 | cycloalkane | C1CCCC1 |
| 138 | alkane | CCC(C)(C(C)C)C(C)(C)C |
| 139 | alcohol | CC(C)C(C)(C)O |
| 140 | alkane | CCCCCCC(C)CC |
| 141 | alcohol | CC[C@H](C)O |
| 142 | alkane | CCCCCCCCCC |

Continued on next page

Table SI 1 – Continued from previous page

| NO. | SMILES | Category |
|---|---|---|
| 143 | alkane | CC[C@@H](C)[C@@H](C)C(C)C |
| 144 | ketone | CCCCC(C)=O |
| 145 | ketone | CCCCC(=O)CCC |

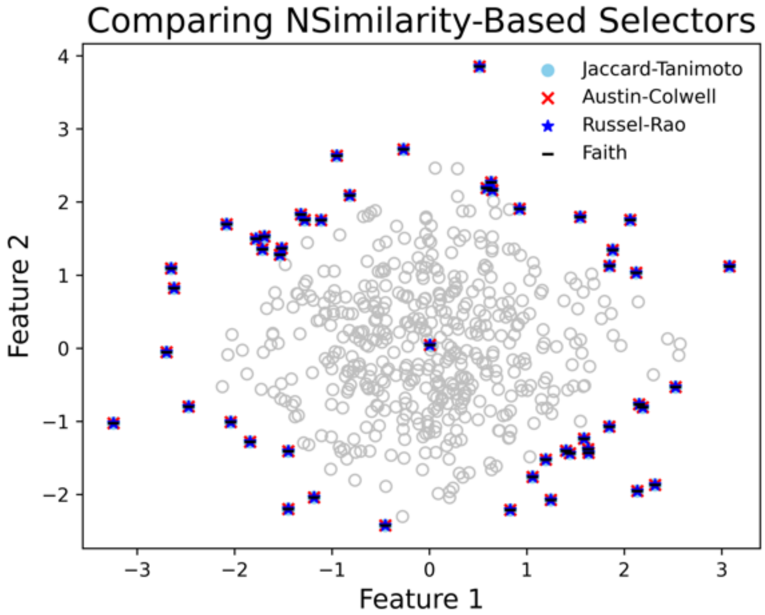


Fig. SI 2 Comparing NSimilarity based selection methods.

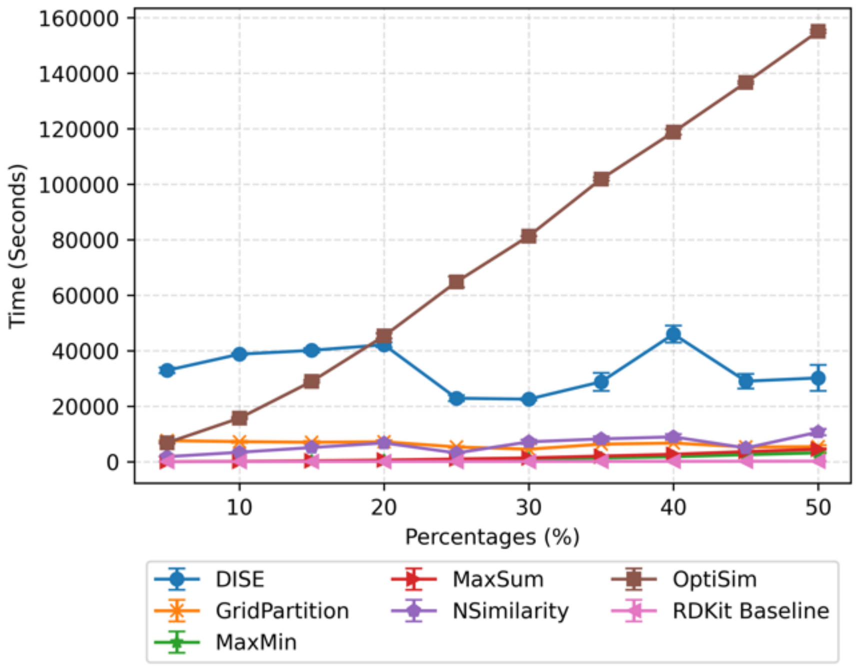


Fig. SI 3 Running time comparisons of different subset selection algorithms. The 1024-bit ECFP4 fingerprints were computed with `RDKit` for the 14k logP dataset from EPA public data[1]. The `MaxMinPicker` method in `RDKit` was as the base line. It is noticed OptiSim are slower because the setup of searching the optimal radius in `Selector` in order to get the required number of selected data points.

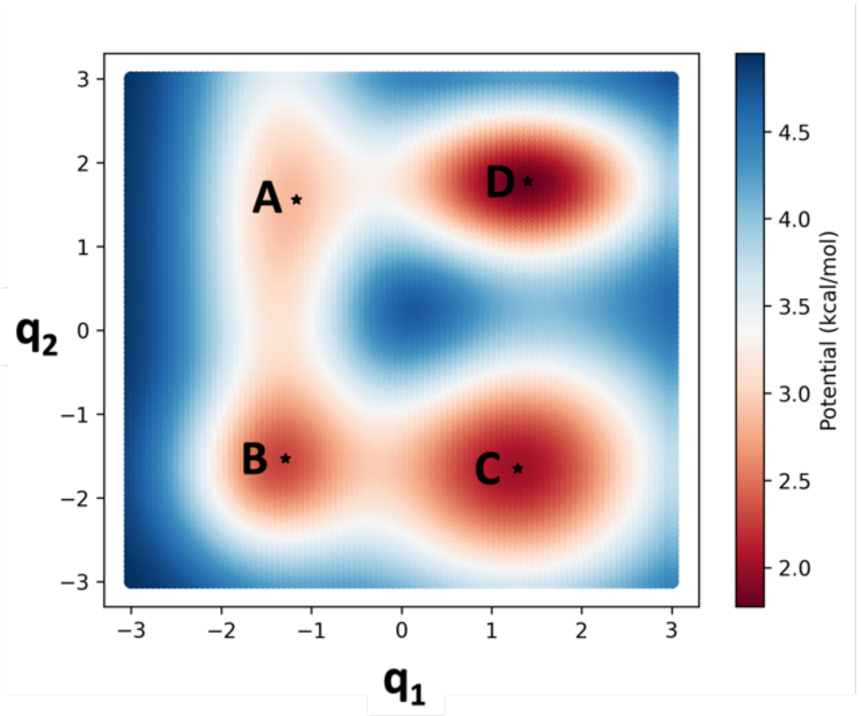


Fig. SI 4 The four-well potential with four minima: $V_A(-1.17, 1.56) = 2.84$, $V_B(-1.29, -1.53) = 2.28$, $V_C(1.29, -1.65) = 2.01$, and $V_D(1.4, 1.78) = 1.77$ kcal/mol.[2]

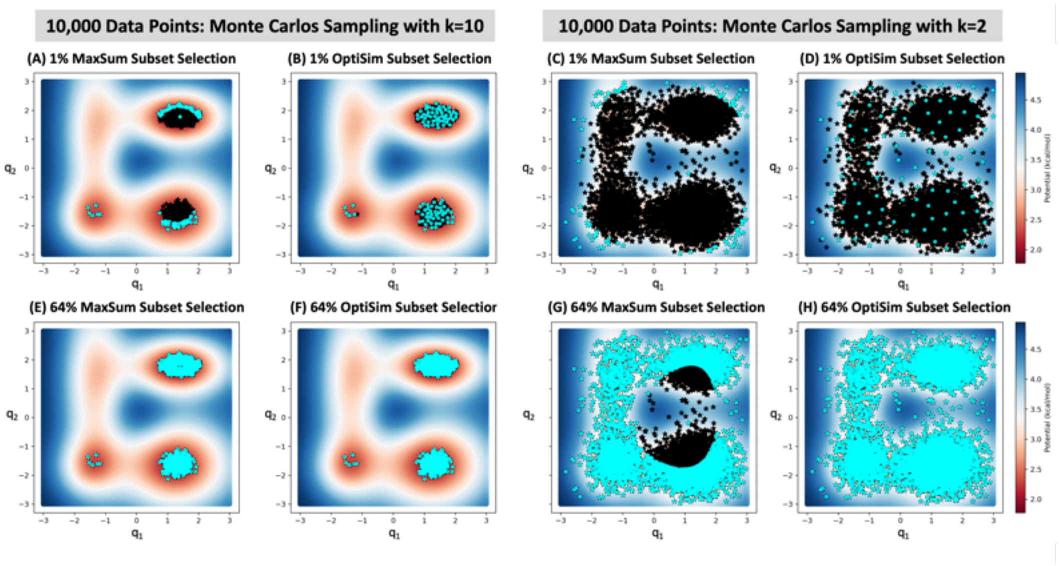


Fig. SI 5 Selecting subsets (colored with cyan circles) of four-well potential with MaxSum (1st and 3rd columns) and OptiSim (2nd and 4th columns) methods from a dataset of 10,000 points (colored with black stars) generated with Monte Carlo sampling using values of k=10 (two left columns) and k=2 (two right columns).

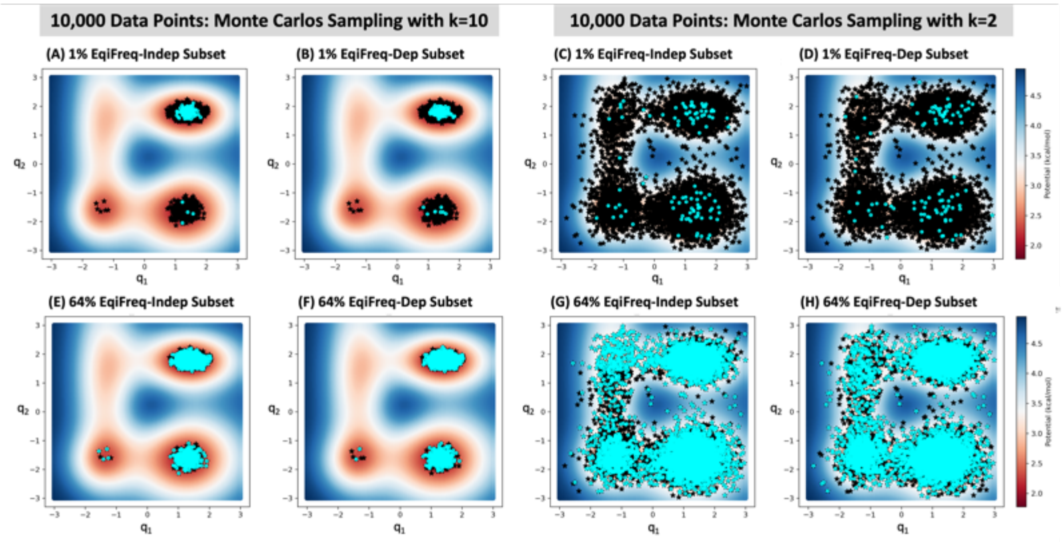


Fig. SI 6 Selecting subsets (colored with cyan circles) of four-well potential with Grid Partition Equifrequent Dependent (1st and 3rd columns) and Independent (2nd and 4th columns) methods from a dataset of 10,000 points (colored with black stars) generated with Monte Carlo sampling using values of k=10 (two left columns) and k=2 (two right columns). For the Grid Partition, 5 bins are used along each access.

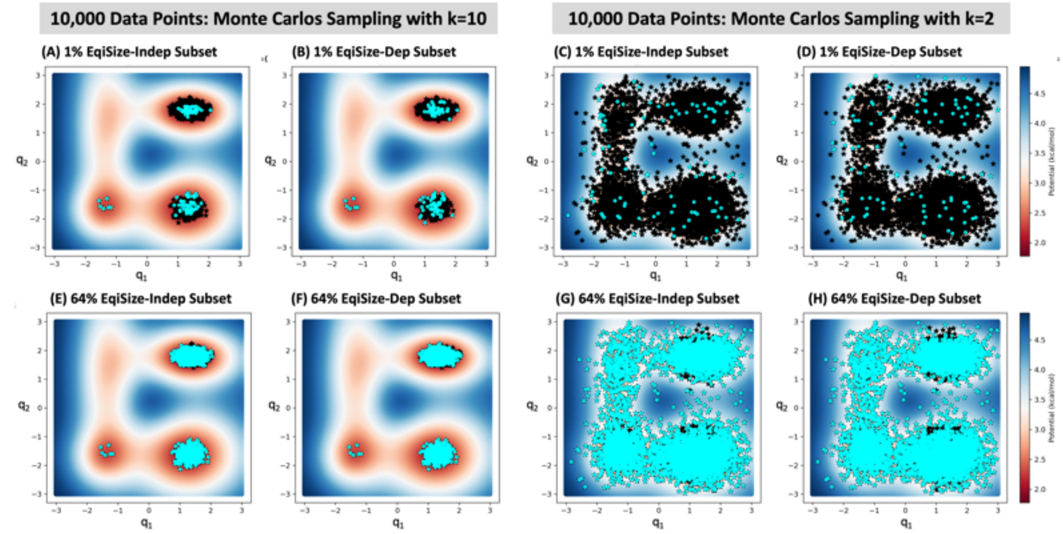


Fig. SI 7 Selecting subsets (colored with cyan circles) of four-well potential with Grid Partition Equisize Dependent (1st and 3rd columns) and Independent (2nd and 4th columns) methods from a dataset of 10,000 points (colored with black stars) generated with Monte Carlo sampling using values of k=10 (two left columns) and k=2 (two right columns). For the Grid Partition, 5 bins are used along each access.

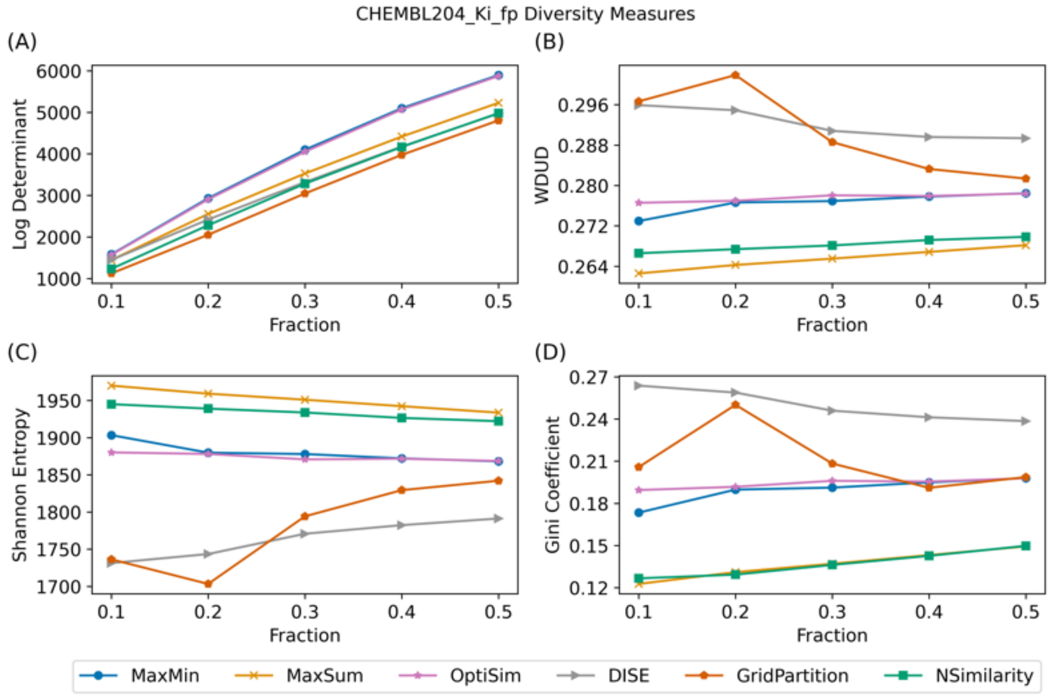


Fig. SI 8 Diversity selection measurements of ChEMBL204 Ki subset ($n_{total} = 2754$) selection with different selection methods. (A) Log-determinant of the selected subset. (B) Wasserstein distance to uniform distribution (WDUD) of the selected subset. (C) Shannon entropy of the selected subset. (D) Gini coefficient of the selected subset. Based on the WDUD, MaxSum and NSimilarity perform the best across the different fractions. Grid partitioning has the lowest diversity when less than 30% of the data is selected, but for larger fractions of the data DISE performs worst. The Gini coefficient, MaxSum and NSimilarity consistently yield the most diverse sets and DISE consistently gives the least diverse set.

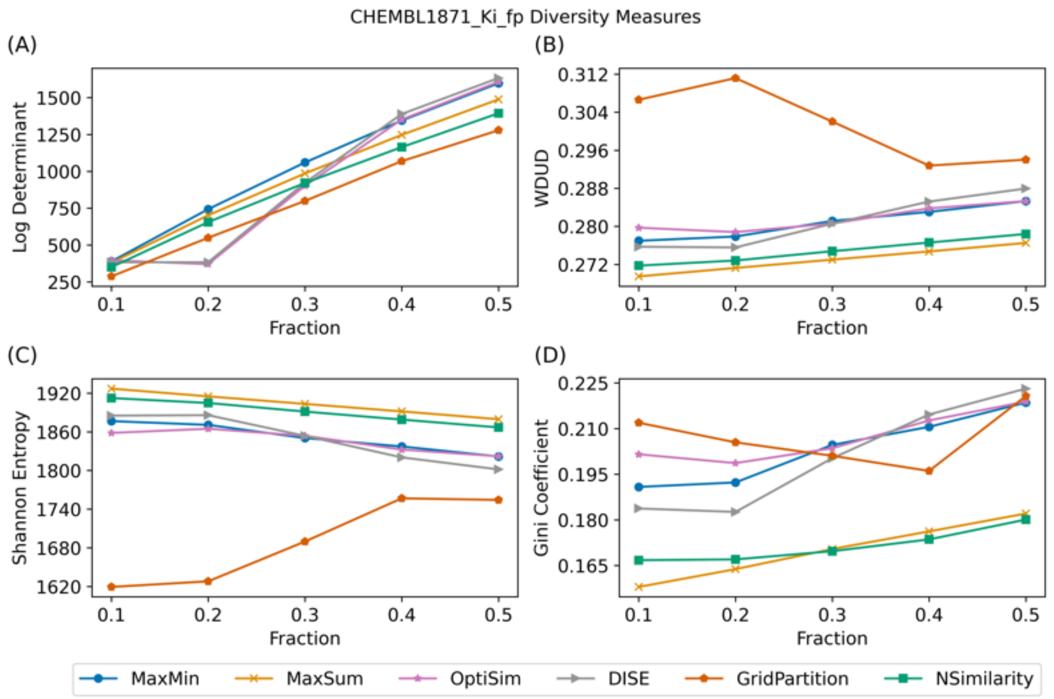


Fig. SI 9 Diversity selection measurements of ChEMBL1871 Ki subset ($n_{total} = 659$) selection with different selection methods. (A) Log-determinant of the selected subset. (B) Wasserstein distance to uniform distribution (WDUD) of the selected subset. (C) Shannon entropy of the selected subset. (D) Gini coefficient of the selected subset.

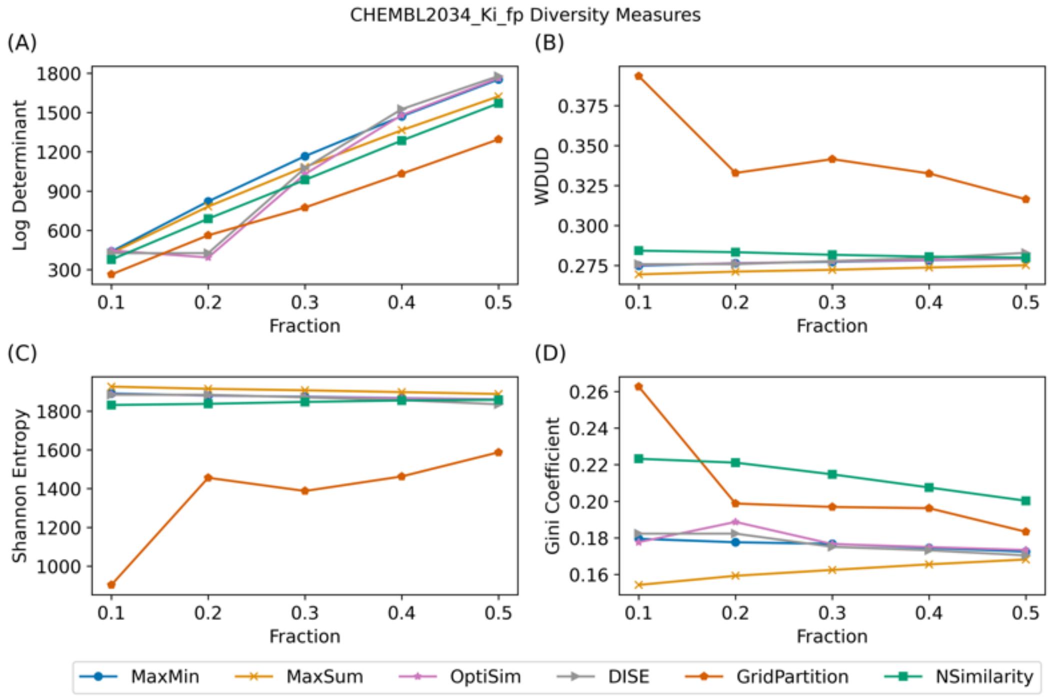


Fig. SI 10 Diversity selection measurements of ChEMBL2034 Ki subset ($n_{total} = 750$) selection with different selection methods. (A) Log-determinant of the selected subset. (B) Wasserstein distance to uniform distribution (WDUD) of the selected subset. (C) Shannon entropy of the selected subset. (D) Gini coefficient of the selected subset.

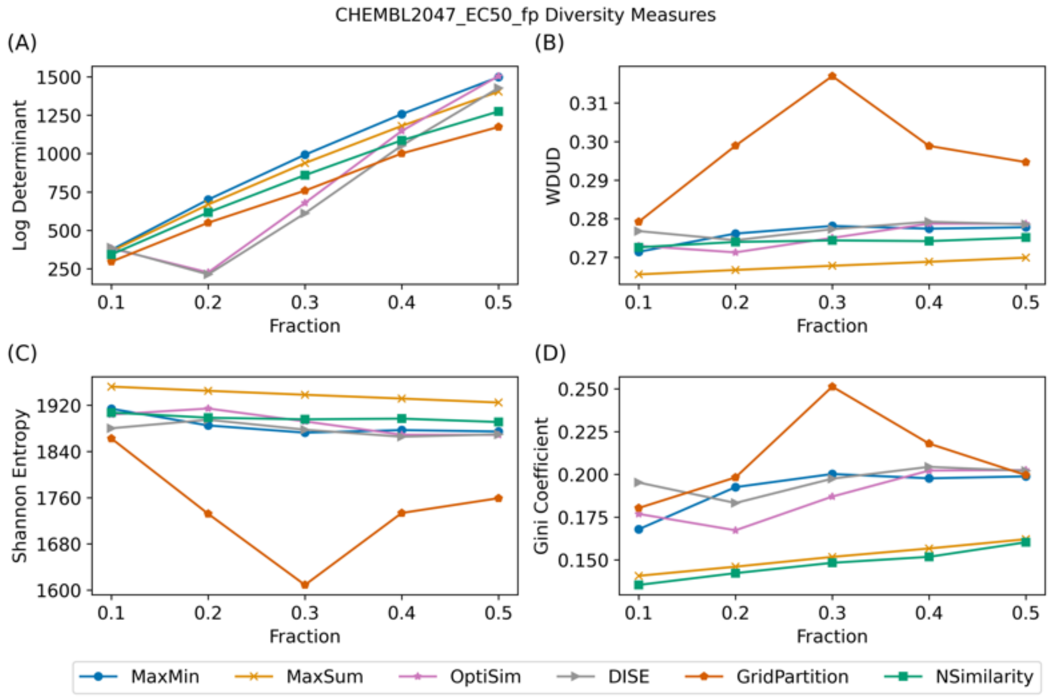


Fig. SI 11 Diversity selection measurements of ChEMBL2047 EC50 subset ($n_{total} = 631$) selection with different selection methods. (A) Log-determinant of the selected subset. (B) Wasserstein distance to uniform distribution (WDUD) of the selected subset. (C) Shannon entropy of the selected subset. (D) Gini coefficient of the selected subset.

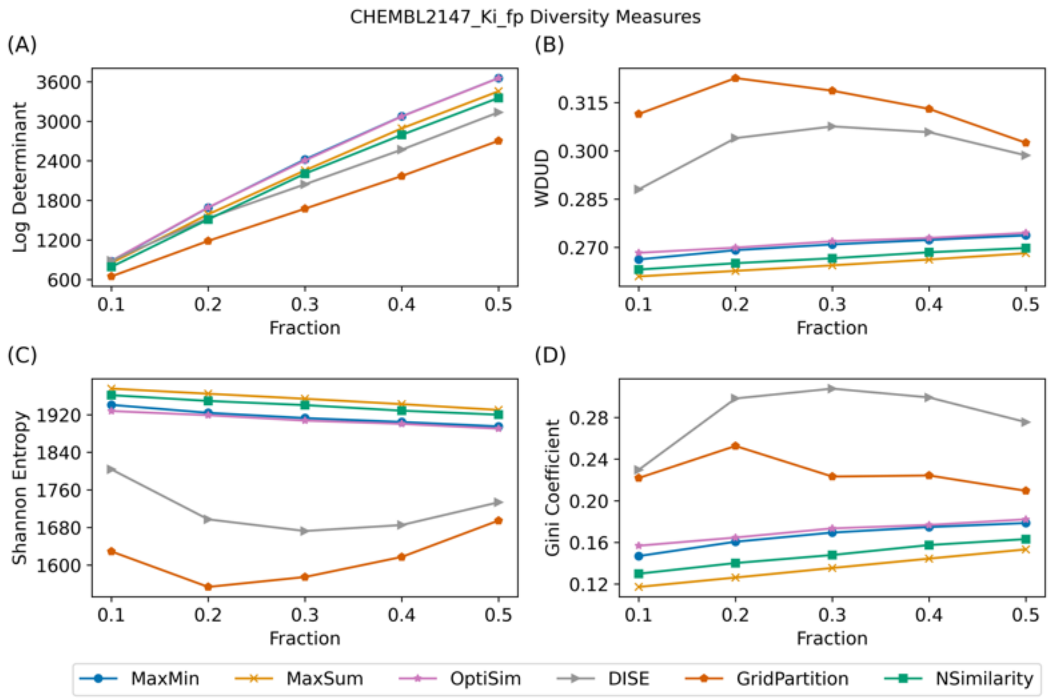


Fig. SI 12 Diversity selection measurements of ChEMBL2147 Ki subset ($n_{total} = 1456$) selection with different selection methods. (A) Log-determinant of the selected subset. (B) Wasserstein distance to uniform distribution (WDUD) of the selected subset. (C) Shannon entropy of the selected subset. (D) Gini coefficient of the selected subset.

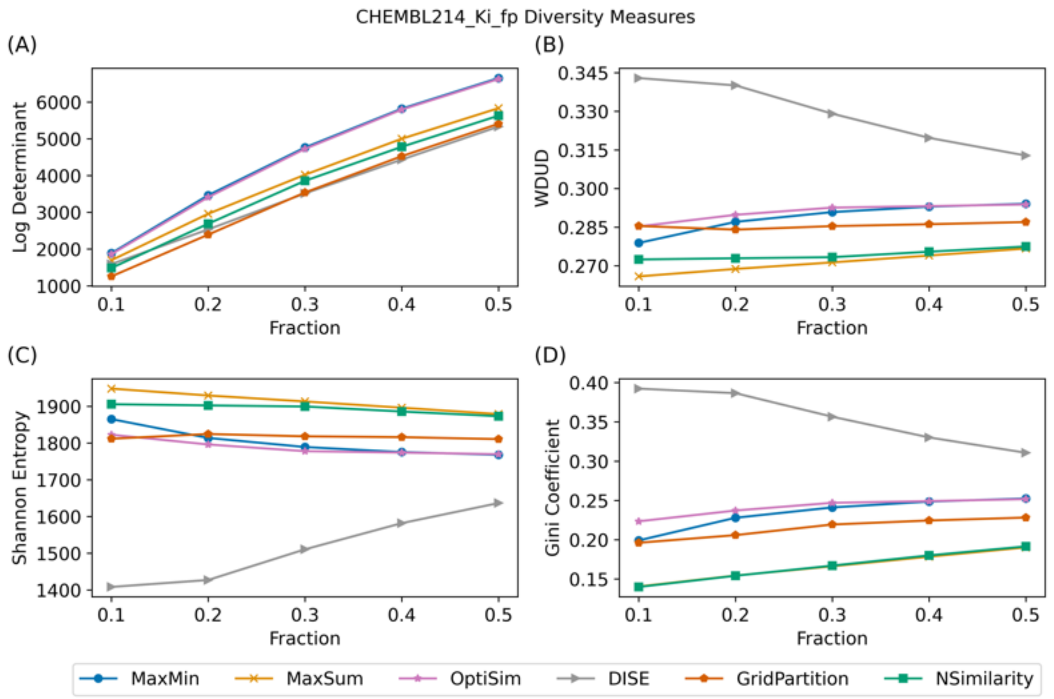


Fig. SI 13 Diversity selection measurements of ChEMBL214 Ki subset ($n_{total} = 3317$) selection with different selection methods. (A) Log-determinant of the selected subset. (B) Wasserstein distance to uniform distribution (WDUD) of the selected subset. (C) Shannon entropy of the selected subset. (D) Gini coefficient of the selected subset.

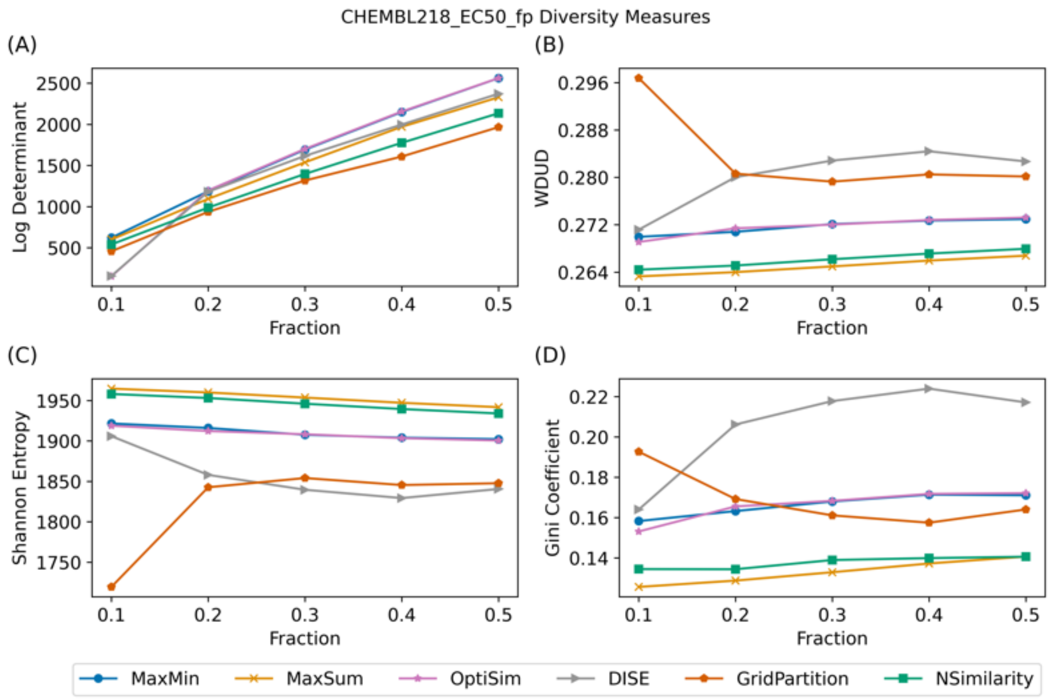


Fig. SI 14 Diversity selection measurements of ChEMBL218 EC50 subset ($n_{total} = 1031$) selection with different selection methods. (A) Log-determinant of the selected subset. (B) Wasserstein distance to uniform distribution (WDUD) of the selected subset. (C) Shannon entropy of the selected subset. (D) Gini coefficient of the selected subset.

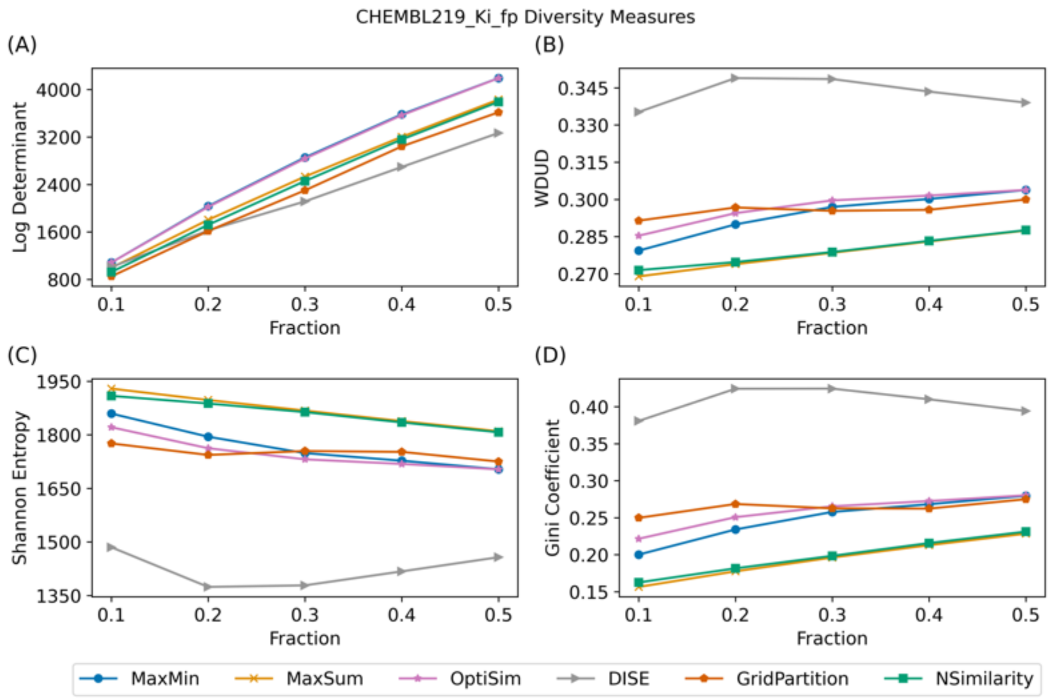


Fig. SI 15 Diversity selection measurements of ChEMBL219 Ki subset ($n_{total} = 1859$) selection with different selection methods. (A) Log-determinant of the selected subset. (B) Wasserstein distance to uniform distribution (WDUD) of the selected subset. (C) Shannon entropy of the selected subset. (D) Gini coefficient of the selected subset.

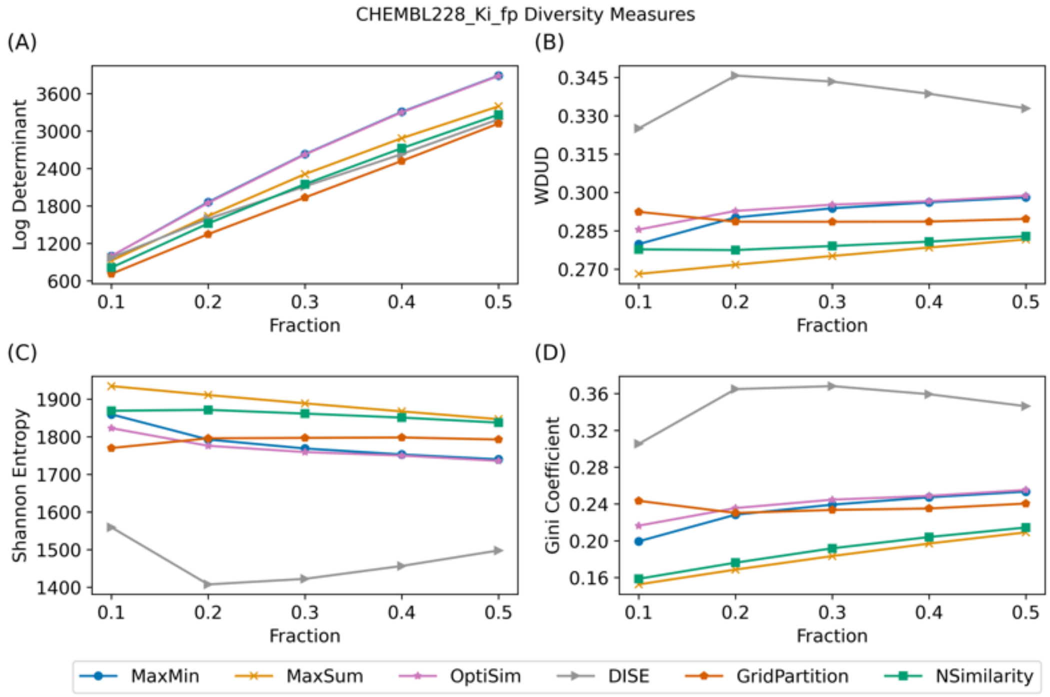


Fig. SI 16 Diversity selection measurements of ChEMBL228 Ki subset ($n_{total} = 1704$) selection with different selection methods.

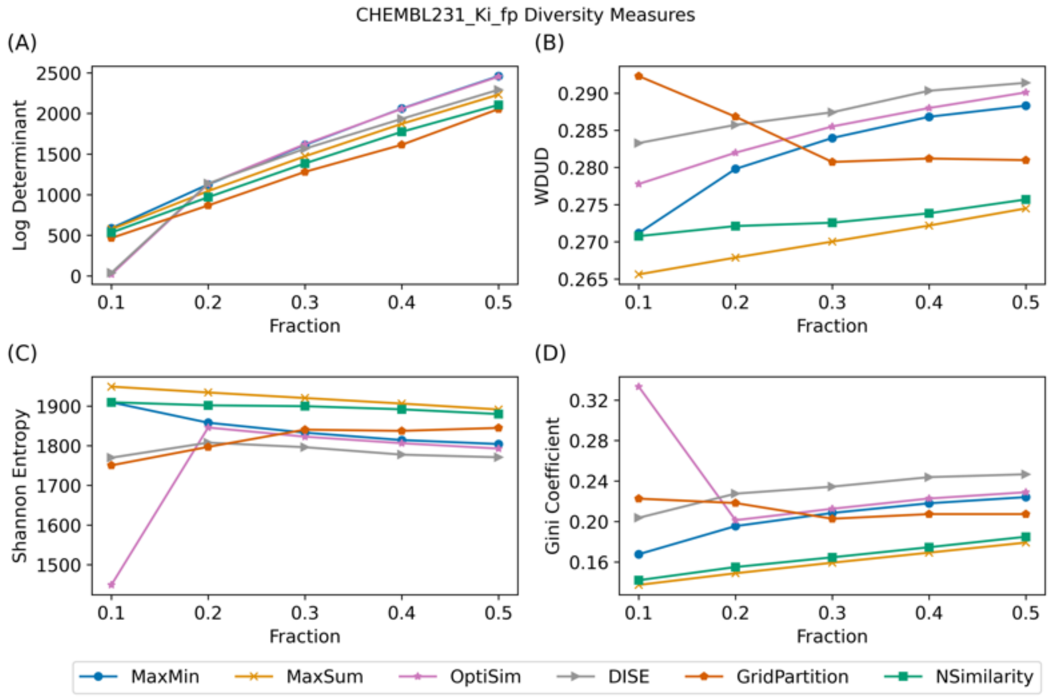


Fig. SI 17 Diversity selection measurements of ChEMBL231 Ki subset ($n_{total} = 973$) selection with different selection methods. (A) Log-determinant of the selected subset. (B) Wasserstein distance to uniform distribution (WDUD) of the selected subset. (C) Shannon entropy of the selected subset. (D) Gini coefficient of the selected subset.

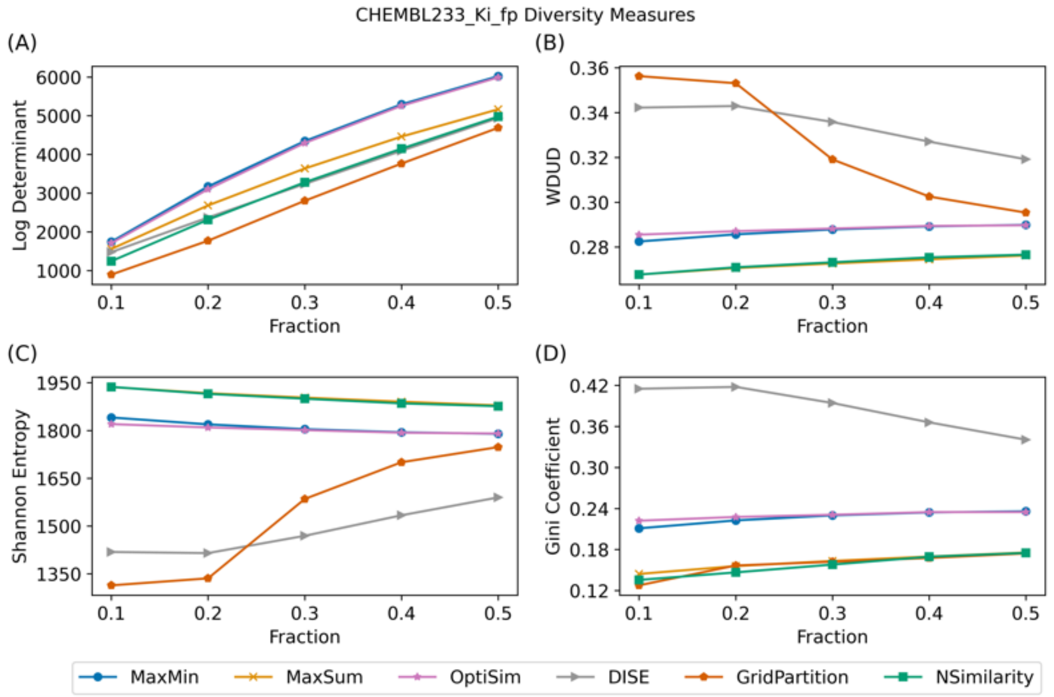


Fig. SI 18 Diversity selection measurements of ChEMBL233 Ki subset ($n_{total} = 3142$) selection with different selection methods. (A) Log-determinant of the selected subset. (B) Wasserstein distance to uniform distribution (WDUD) of the selected subset. (C) Shannon entropy of the selected subset. (D) Gini coefficient of the selected subset.

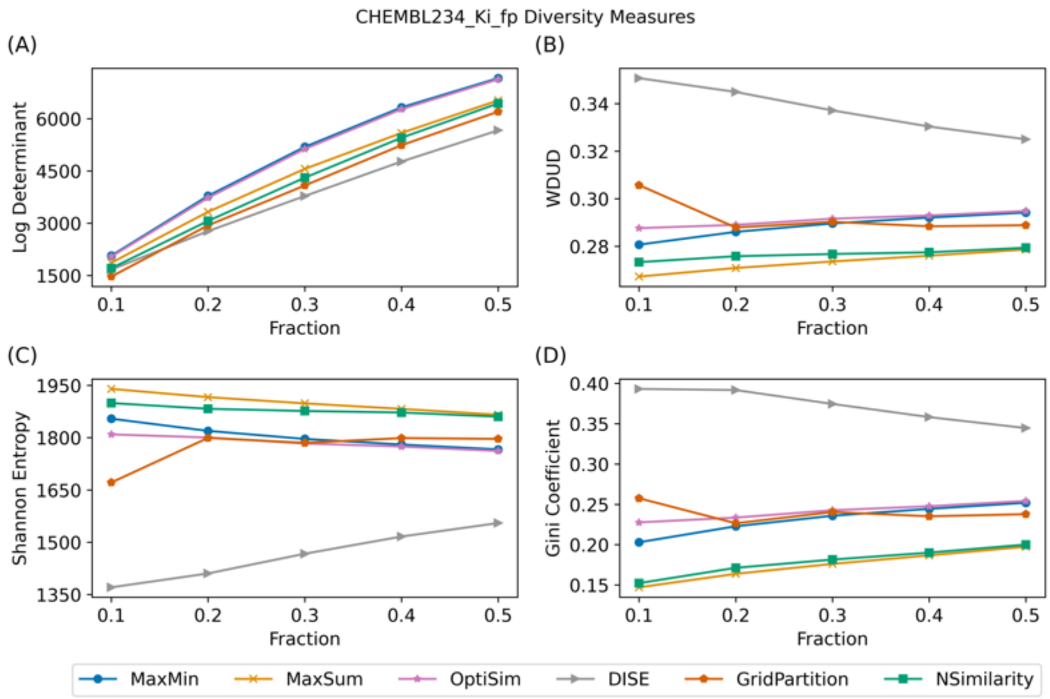


Fig. SI 19 Diversity selection measurements of ChEMBL234 Ki subset ($n_{total} = 3657$) selection with different selection methods. (A) Log-determinant of the selected subset. (B) Wasserstein distance to uniform distribution (WDUD) of the selected subset. (C) Shannon entropy of the selected subset. (D) Gini coefficient of the selected subset.

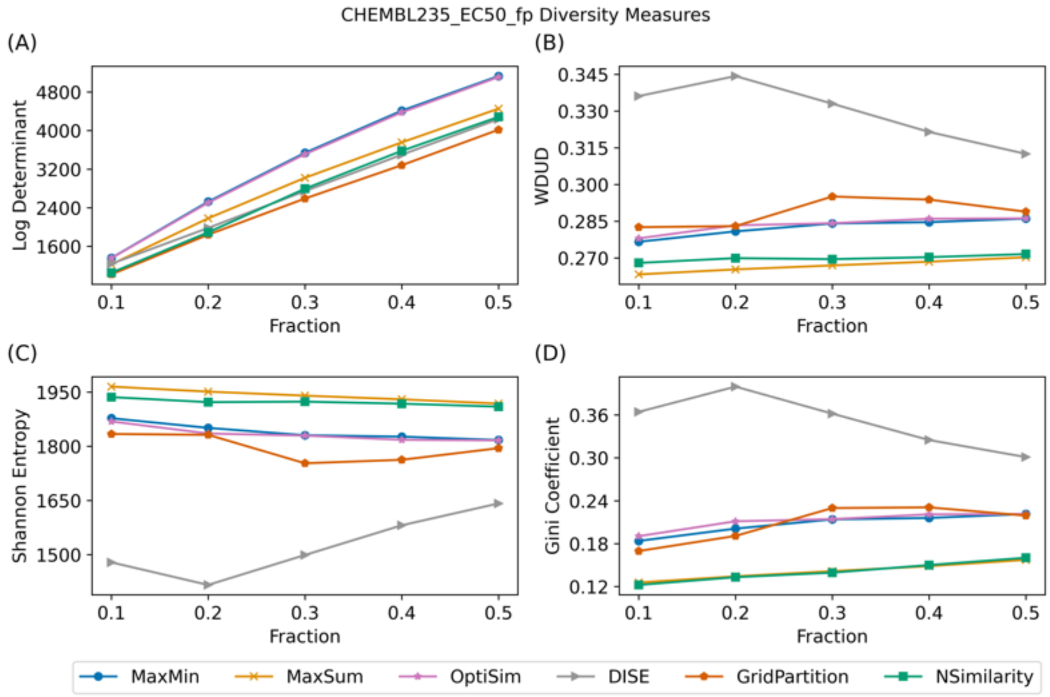


Fig. SI 20 Diversity selection measurements of ChEMBL235 EC50 subset ($n_{total} = 2349$) selection with different selection methods. (A) Log-determinant of the selected subset. (B) Wasserstein distance to uniform distribution (WDUD) of the selected subset. (C) Shannon entropy of the selected subset. (D) Gini coefficient of the selected subset.

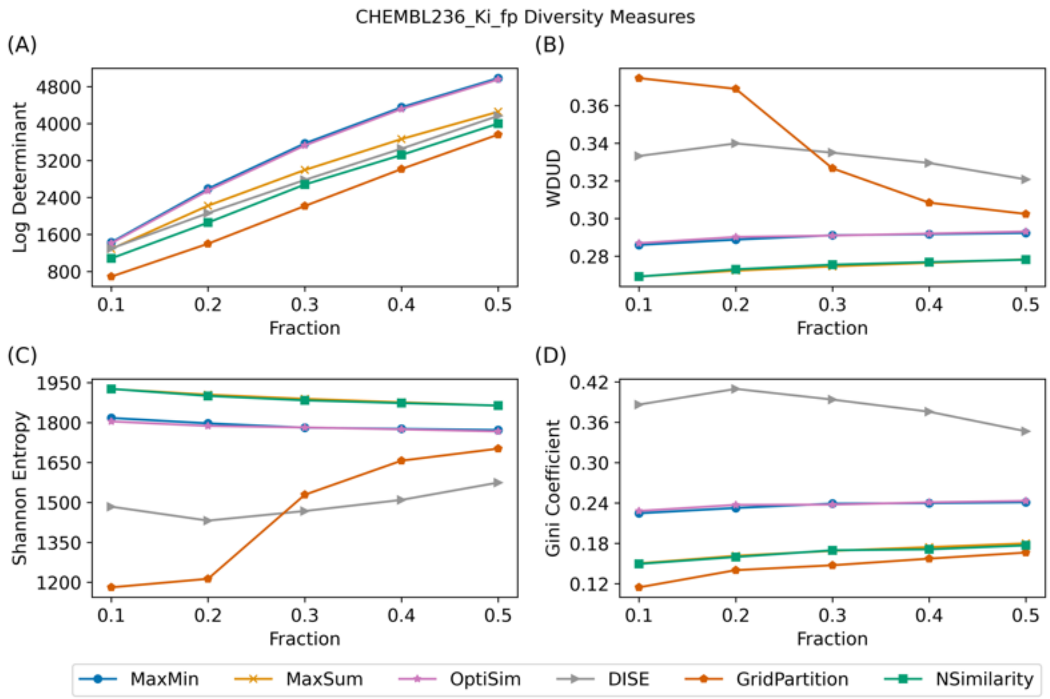


Fig. SI 21 Diversity selection measurements of ChEMBL236 Ki subset ($n_{total} = 2598$) selection with different selection methods. (A) Log-determinant of the selected subset. (B) Wasserstein distance to uniform distribution (WDUD) of the selected subset. (C) Shannon entropy of the selected subset. (D) Gini coefficient of the selected subset.

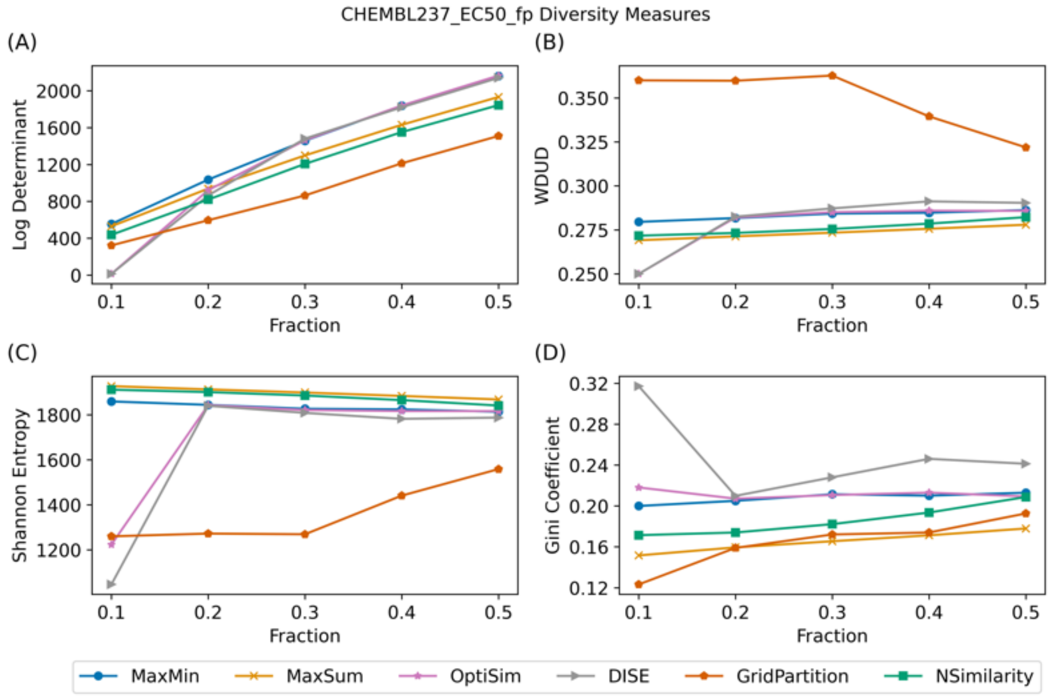


Fig. SI 22 Diversity selection measurements of ChEMBL237 EC50 subset ($n_{total} = 955$) selection with different selection methods.

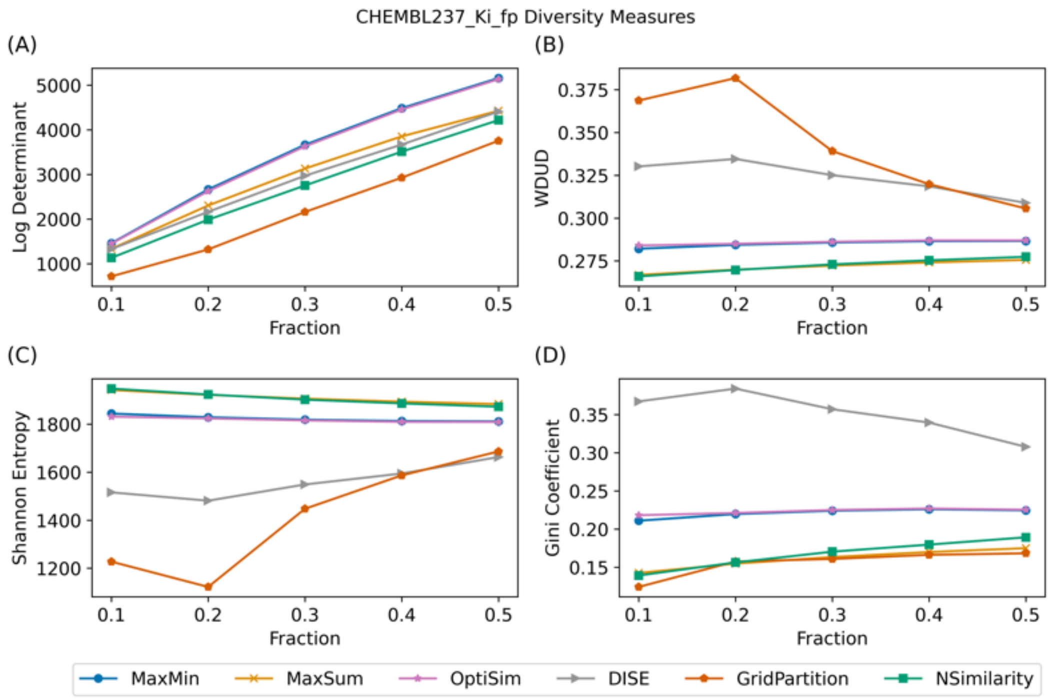


Fig. SI 23 Diversity selection measurements of ChEMBL237 Ki subset ($n_{total} = 2602$) selection with different selection methods. (A) Log-determinant of the selected subset. (B) Wasserstein distance to uniform distribution (WDUD) of the selected subset. (C) Shannon entropy of the selected subset. (D) Gini coefficient of the selected subset.

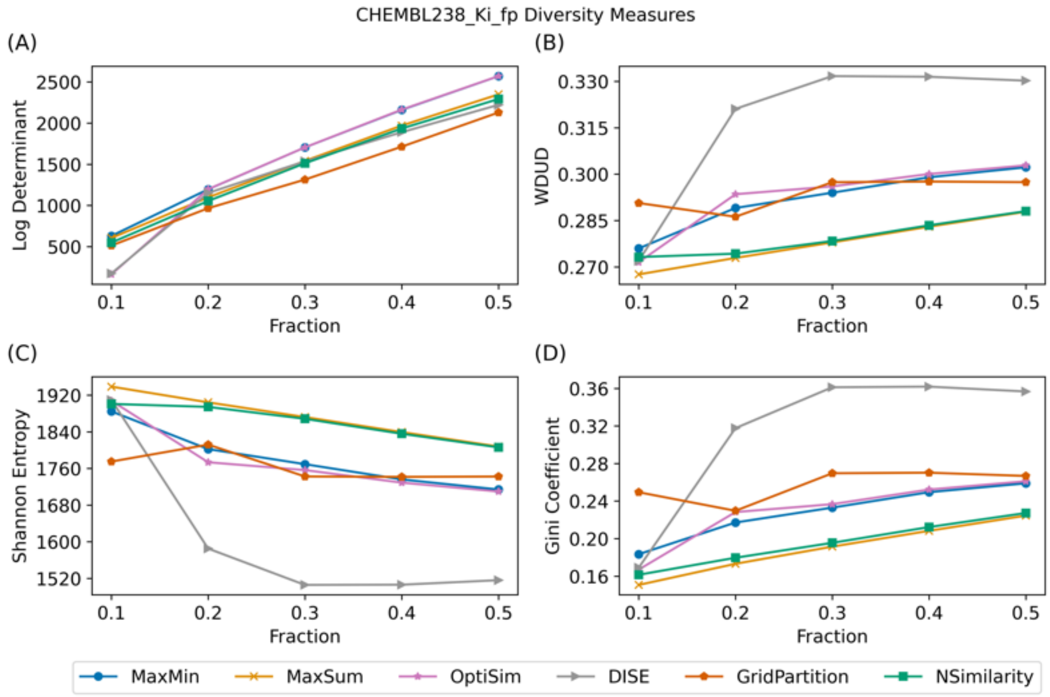


Fig. SI 24 Diversity selection measurements of ChEMBL238 Ki subset ($n_{total} = 1052$) selection with different selection methods. (A) Log-determinant of the selected subset. (B) Wasserstein distance to uniform distribution (WDUD) of the selected subset. (C) Shannon entropy of the selected subset. (D) Gini coefficient of the selected subset.

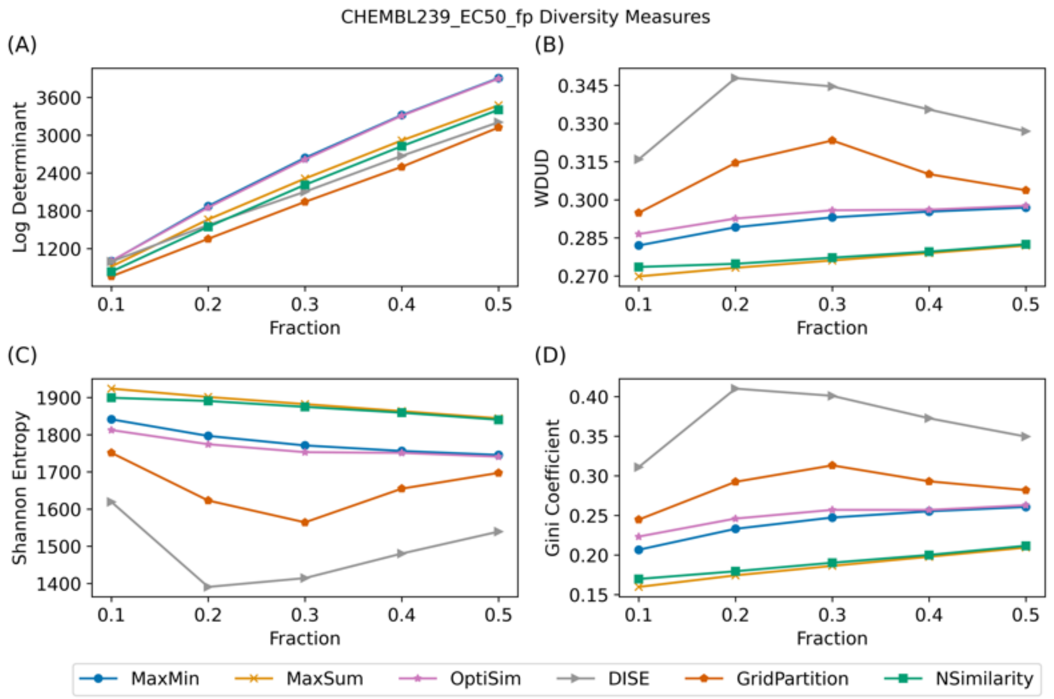


Fig. SI 25 Diversity selection measurements of ChEMBL239 EC50 subset ($n_{total} = 1721$) selection with different selection methods. (A) Log-determinant of the selected subset. (B) Wasserstein distance to uniform distribution (WDUD) of the selected subset. (C) Shannon entropy of the selected subset. (D) Gini coefficient of the selected subset.

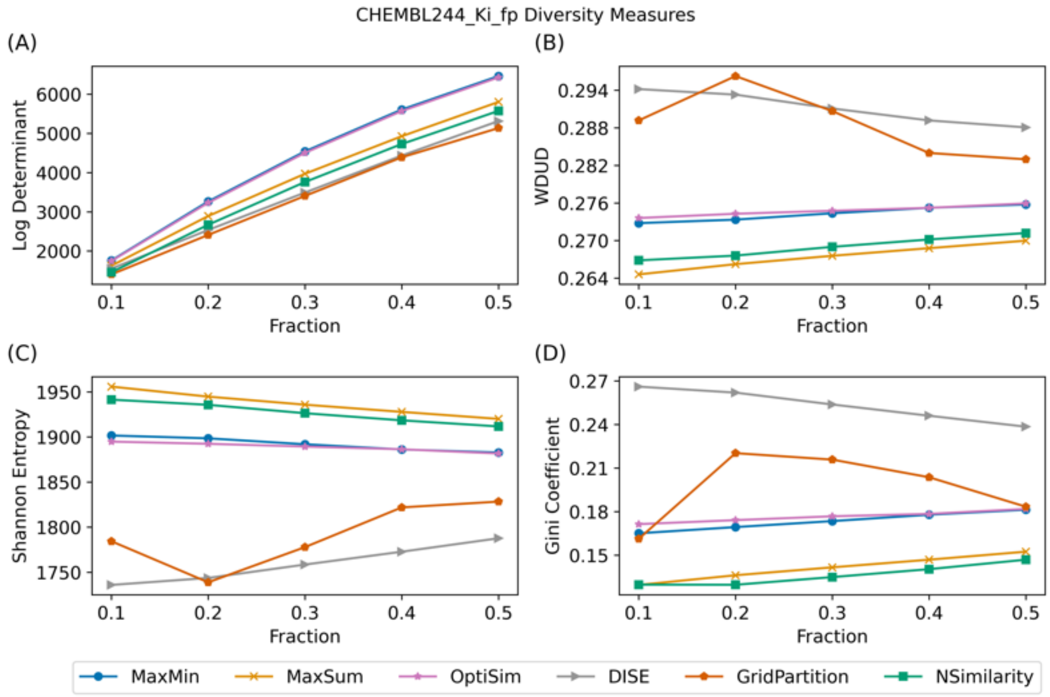


Fig. SI 26 Diversity selection measurements of ChEMBL244 Ki subset ($n_{total} = 3097$) selection with different selection methods. (A) Log-determinant of the selected subset. (B) Wasserstein distance to uniform distribution (WDUD) of the selected subset. (C) Shannon entropy of the selected subset. (D) Gini coefficient of the selected subset.

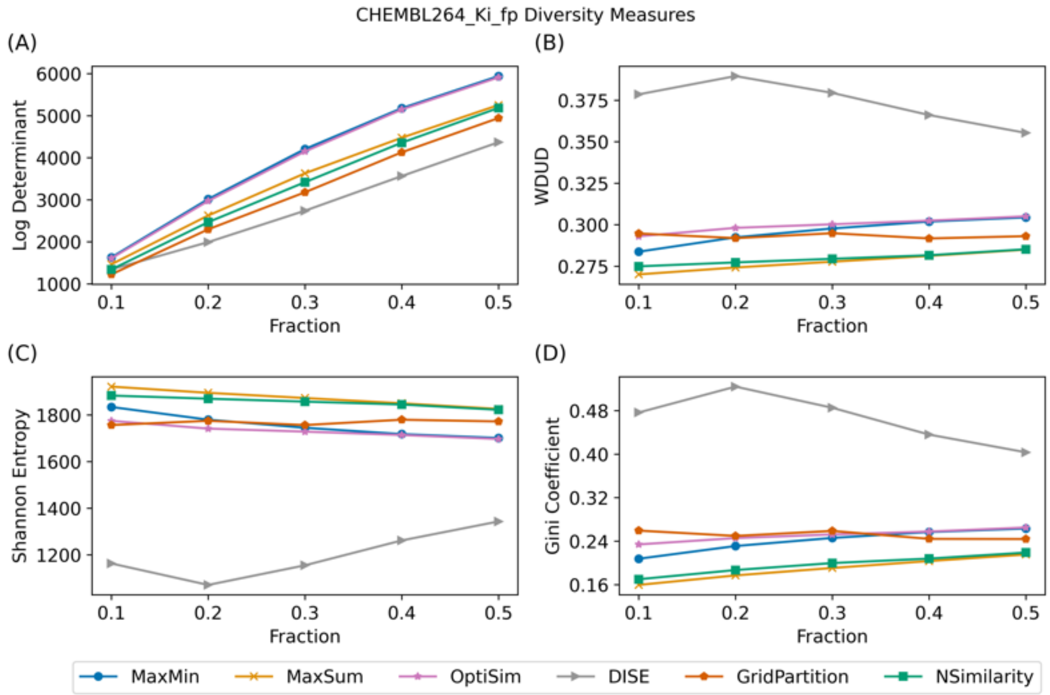


Fig. SI 27 Diversity selection measurements of ChEMBL264 Ki subset ($n_{total} = 2862$) selection with different selection methods. (A) Log-determinant of the selected subset. (B) Wasserstein distance to uniform distribution (WDUD) of the selected subset. (C) Shannon entropy of the selected subset. (D) Gini coefficient of the selected subset.

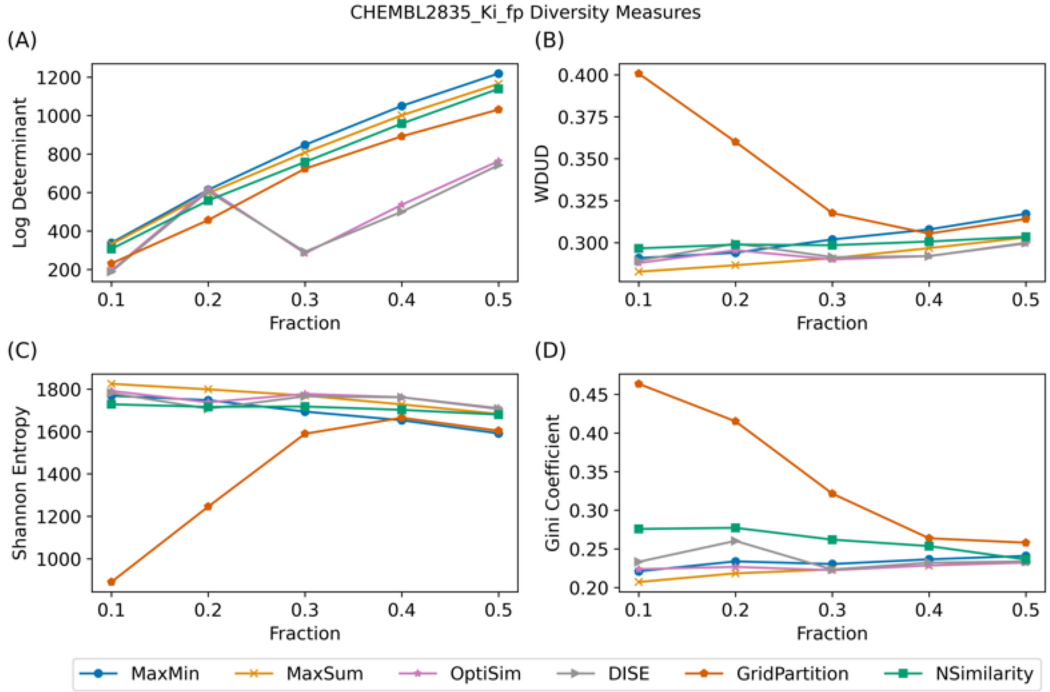


Fig. SI 28 Diversity selection measurements of ChEMBL2835 Ki subset ($n_{total} = 615$) selection with different selection methods. (A) Log-determinant of the selected subset. (B) Wasserstein distance to uniform distribution (WDUD) of the selected subset. (C) Shannon entropy of the selected subset. (D) Gini coefficient of the selected subset.

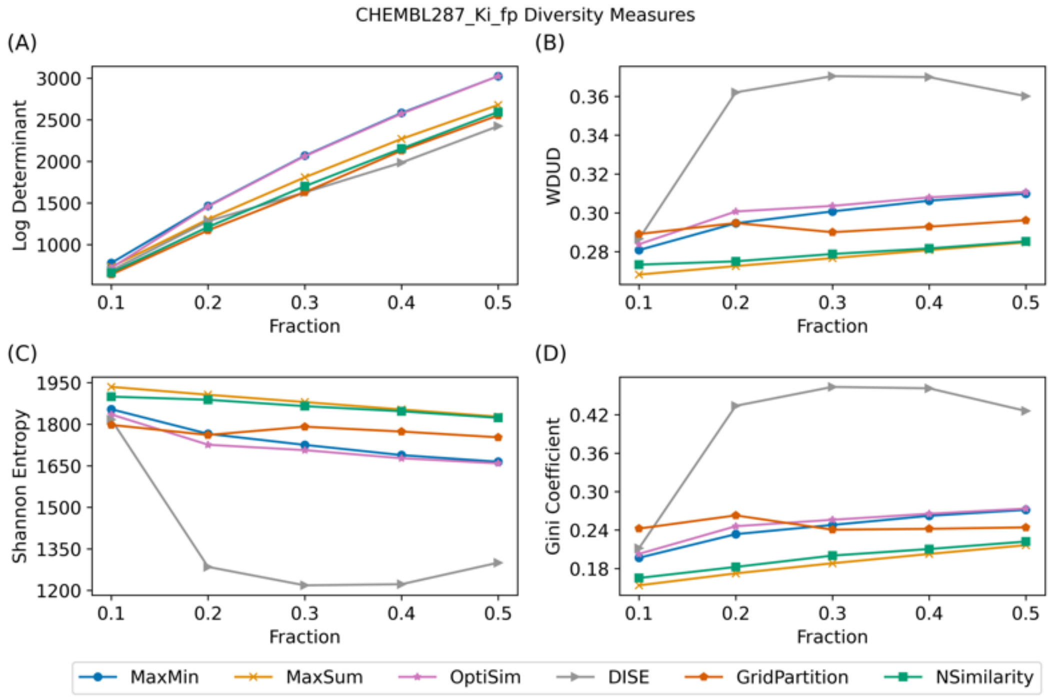


Fig. SI 29 Diversity selection measurements of ChEMBL287 Ki subset ($n_{total} = 1328$) selection with different selection methods. (A) Log-determinant of the selected subset. (B) Wasserstein distance to uniform distribution (WDUD) of the selected subset. (C) Shannon entropy of the selected subset. (D) Gini coefficient of the selected subset.

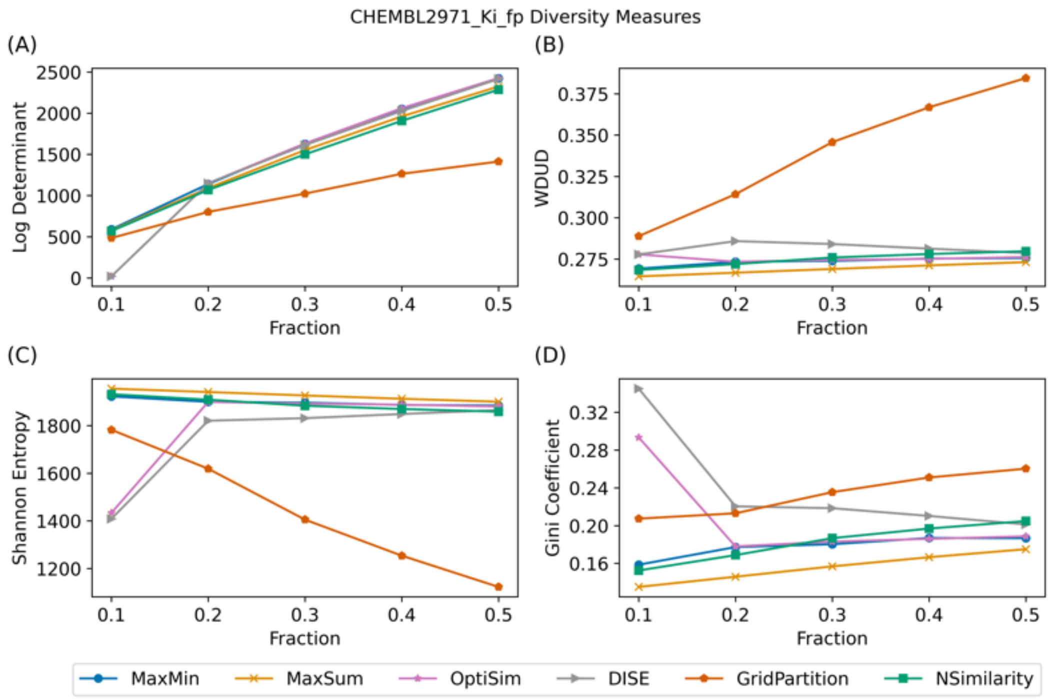


Fig. SI 30 Diversity selection measurements of ChEMBL2971 Ki subset ($n_{total} = 976$) selection with different selection methods. (A) Log-determinant of the selected subset. (B) Wasserstein distance to uniform distribution (WDUD) of the selected subset. (C) Shannon entropy of the selected subset. (D) Gini coefficient of the selected subset.

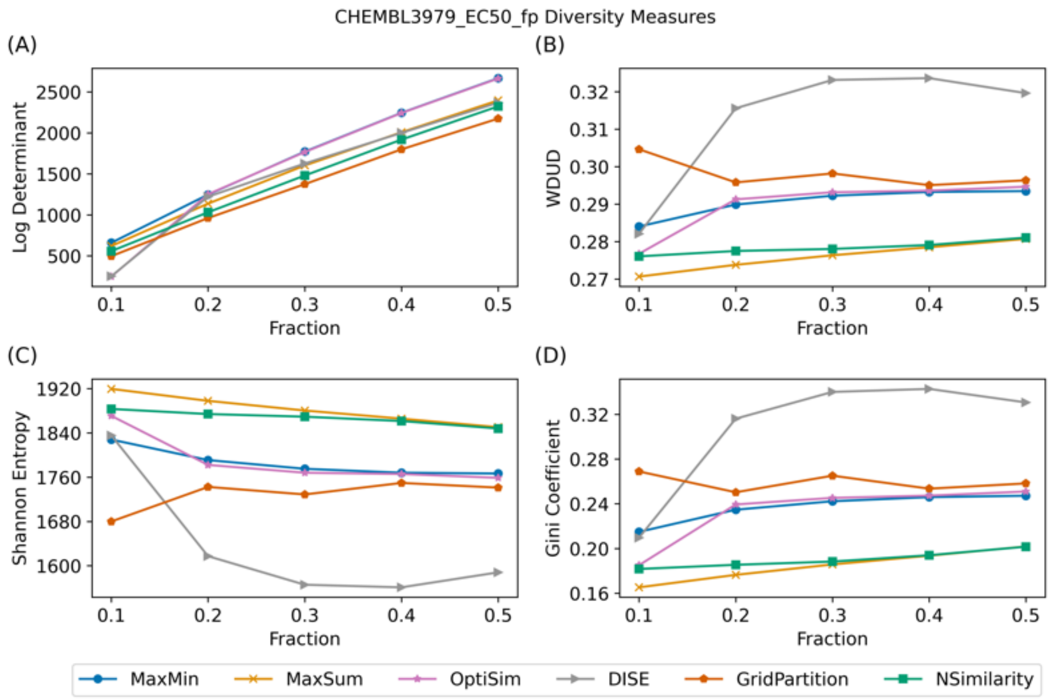


Fig. SI 31 Diversity selection measurements of ChEMBL3979 EC50 subset ($n_{total} = 1125$) selection with different selection methods. (A) Log-determinant of the selected subset. (B) Wasserstein distance to uniform distribution (WDUD) of the selected subset. (C) Shannon entropy of the selected subset. (D) Gini coefficient of the selected subset.

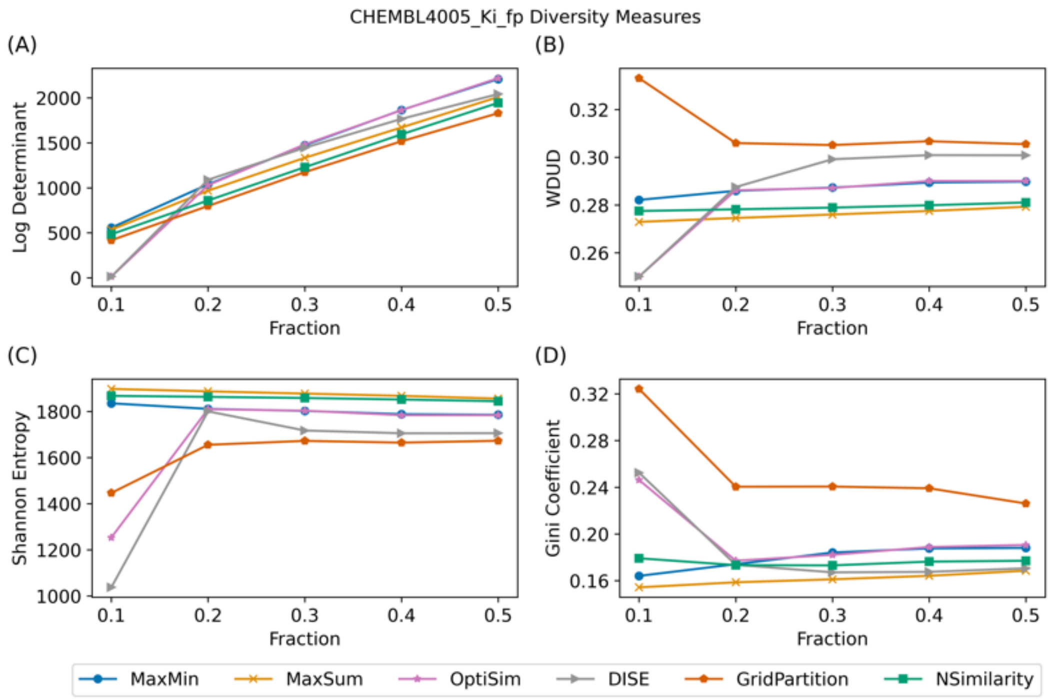


Fig. SI 32 Diversity selection measurements of ChEMBL4005 Ki subset ($n_{total} = 960$) selection with different selection methods. (A) Log-determinant of the selected subset. (B) Wasserstein distance to uniform distribution (WDUD) of the selected subset. (C) Shannon entropy of the selected subset. (D) Gini coefficient of the selected subset.

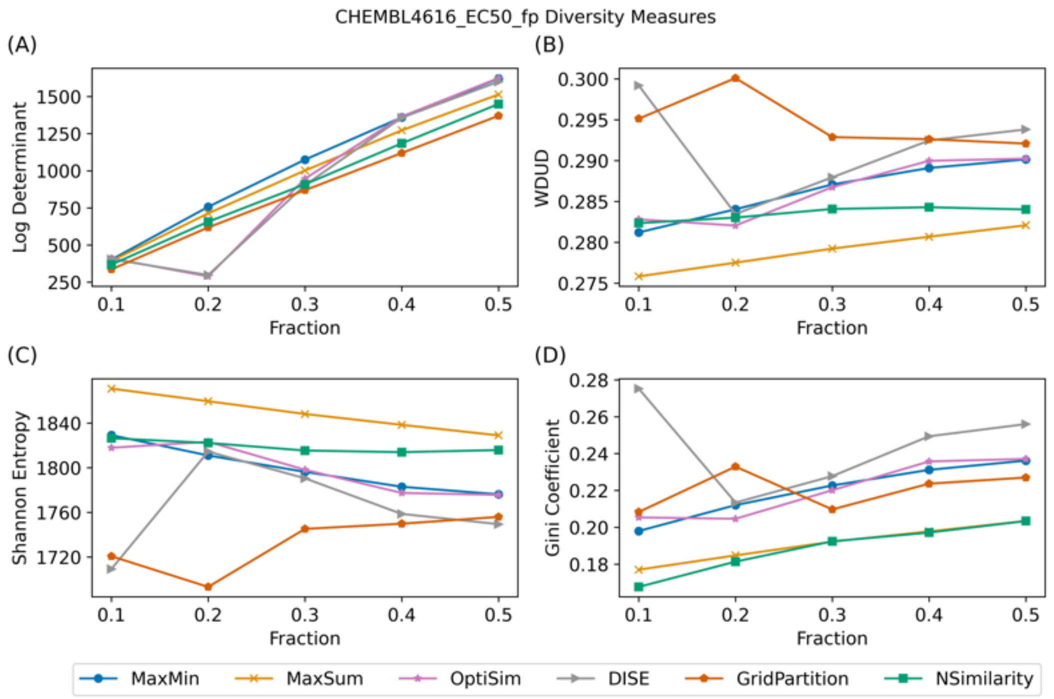


Fig. SI 33 Diversity selection measurements of ChEMBL4616 EC50 subset ($n_{total} = 682$) selection with different selection methods. (A) Log-determinant of the selected subset. (B) Wasserstein distance to uniform distribution (WDUD) of the selected subset. (C) Shannon entropy of the selected subset. (D) Gini coefficient of the selected subset.

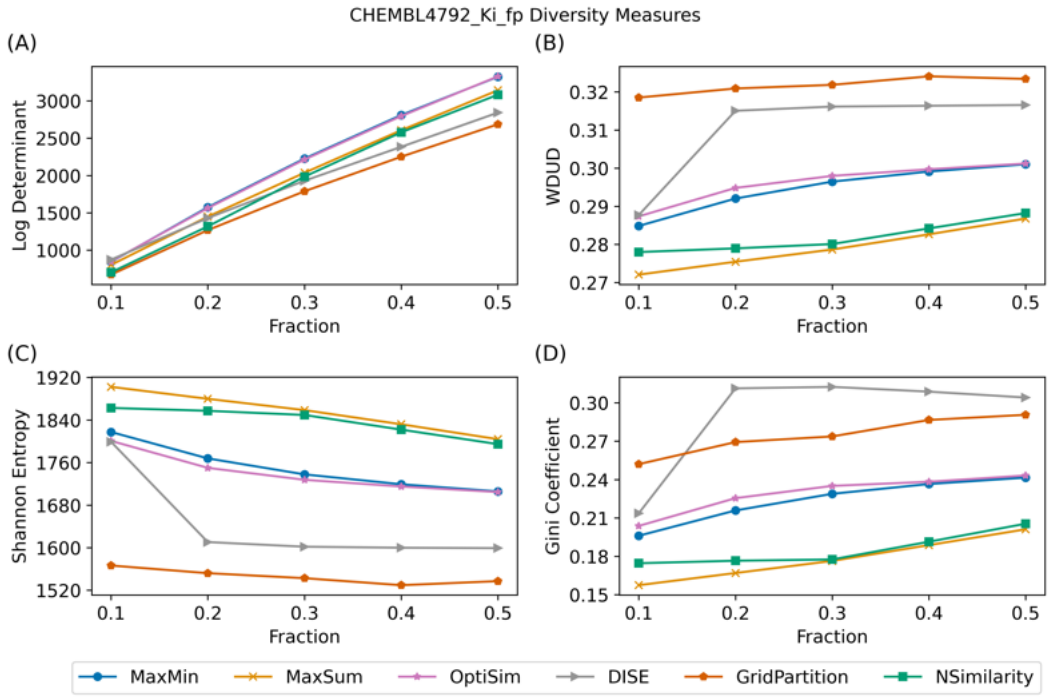


Fig. SI 34 Diversity selection measurements of ChEMBL4792 Ki subset ($n_{total} = 1471$) selection with different selection methods. (A) Log-determinant of the selected subset. (B) Wasserstein distance to uniform distribution (WDUD) of the selected subset. (C) Shannon entropy of the selected subset. (D) Gini coefficient of the selected subset.
